# Polymorphisms in *Brucella* Carbonic anhydrase II mediate CO_2_ dependence and fitness *in vivo*

**DOI:** 10.1101/804740

**Authors:** JM García-Lobo, Y Ortiz, C González-Riancho, A Seoane, B Arellano-Reynoso, FJ Sangari

**Author notes:** FJS conceived and coordinated the study, conducted bacteriology work and wrote the manuscript. JMGL analyzed the data and wrote the manuscript. YO, CGR, AS and BAR conducted bacteriology work. All authors interpreted the data, corrected the manuscript, and approved the content for publication.

## Abstract

Some *Brucella* isolates are known to require an increased concentration of CO_2_ for growth, especially in the case of primary cultures obtained directly from infected animals. Moreover, the different *Brucella* species and biovars show a characteristic pattern of CO_2_ requirement, and this trait has been included among the routine typing tests used for species and biovar differentiation. By comparing the differences in gene content among different CO_2_-dependent and CO_2_-independent *Brucella* strains we have confirmed that carbonic anhydrase II (CA II), is the enzyme responsible for this phenotype in all the *Brucella* strains tested. *Brucella* species contain two carbonic anhydrases of the β family, CA I and CA II; genetic polymorphisms exist for both of them in different isolates, but only those putatively affecting the activity of CA II correlate with the CO_2_ requirement of the corresponding isolate. Analysis of these polymorphisms does not allow the determination of CA I functionality, while the polymorphisms in CA II consist of small deletions that cause a frameshift that changes the C-terminus of the protein, probably affecting its dimerization status, essential for the activity.

CO_2_-independent mutants arise easily *in vitro*, although with a low frequency ranging from 10^−6^ to 10^−10^ depending on the strain. These mutants carry compensatory mutations that produce a full length CA II. At the same time, no change was observed in the sequence coding for CA I. A competitive index assay designed to evaluate the fitness of a CO_2_-dependent strain compared to its corresponding CO_2_-independent strain revealed that while there is no significant difference when the bacteria are grown in culture plates, growth *in vivo* in a mouse model of infection provides a significant advantage to the CO_2_-dependent strain. This could explain why some *Brucella* isolates are CO_2_-dependent in primary isolation. The polymorphism described here also allows the *in silico* determination of the CO_2_ requirement status of any *Brucella* strain.

## Introduction

*Brucella* species are facultative intracellular Gram-negative coccobacilli that cause brucellosis, the most prevalent zoonosis with more than 500,000 human cases reported worldwide every year (Pappas et al., 2006). *Brucella* isolates are routinely identified and classified by biochemical and phenotypical characteristics like urease activity, CO_2_ dependence, H_2_S production, erythritol and dye-sensitivity, lysis by *Brucella*-specific bacteriophages, agglutination with monospecific sera, or even host preference (Alton et al., 1988). The first observations pertaining *Brucella* and CO_2_ were made by Nowak (1908), who noticed that *B. abortus* was more easily isolated from the host tissues when the concentration of oxygen in the atmosphere was reduced, but it was Wilson (1931) who established the requirement of CO_2_ for growth in these isolates. This requirement is not universal within *Brucellaceae*, and the different species and biovars show a characteristic pattern of CO_2_ dependence. Within the classical species, *B. abortus* biovars 1, 2, 3, 4, and some isolates from biovar 9, as well as *B. ovis*, require an increased concentration of CO_2_ for growth, especially in the case of primary cultures obtained directly from infected animals. Within the more recently described species, most strains of *B. pinnipedialis* require supplementary CO_2_ for growth, and most of *B. ceti* do not (Foster et al., 1996). The CO_2_-dependence may be lost by subculturing *in vitro*, with an estimated frequency of 3 x 10^−10^ per cell division (Marr and Wilson, 1950), and this is what happened with well-known laboratory *B. abortus* biovar 1 strains like 2308 or S19, that grow in ambient air.

Facultative intracellular bacteria face two environmental conditions with very dissimilar concentrations of carbon dioxide (CO_2_). Inside mammalian cells, CO_2_ concentration may be as high as 5%, while atmospheric concentration is currently estimated at 0.04%. CO_2_ and bicarbonate (HCO_3_^−^) are essential growth factors for bacteria, and they can be interconverted spontaneously at significant rates. The reversible hydration of CO_2_^−^ into HCO_3_ can also be catalyzed by carbonic anhydrase (CA), a ubiquitous metalloenzyme fundamental to many biological functions including photosynthesis, respiration, and CO_2_ and ion transport. The CA superfamily (CAs, EC 4.2.1.1) has been found in all the three domains of life (Eubacteria, Archaea, and Eukarya) and it currently includes seven known families (α-, β-, γ-, δ-, ζ- η-, and θ-CAs) of distinct evolutionary origin (Supuran, 2018). The conversion of CO_2_ into HCO_3_^−^ is accelerated in the presence of CA and has the effect of ensuring correct CO_2_ concentration for carboxylating enzymes involved in central, amino acid and nucleotide metabolism (Merlin et al., 2003).

CA has been shown to be required to support growth under ambient air in a number of microorganisms like *Ralstonia eutropha* (Kusian et al., 2002), *Escherichia coli* (Hashimoto and Kato, 2003; Merlin et al., 2003), *Corynebacterium glutamicum* (Mitsuhashi et al., 2004), *Saccharomyces cerevisiae* (Aguilera et al., 2005). Growth of CA mutants of these organisms was only possible under an atmosphere with high levels of CO_2_,phenomenon that is explained by the availability of bicarbonate, which is substrate for various carboxylation reactions of physiological importance. These reactions are catalyzed by several housekeeping enzymes. like 5’-phosphoribosyl-5-amino-4-imidazole carboxylase (EC 4.1.1.21), phosphoenolpyruvate carboxylase (EC 4.1.1.31), carbamoyl phosphate synthetase (EC 6.3.4.16), pyruvate carboxylase (EC 6.4.1.1), and acetyl-CoA carboxylase (EC 6.4.1.2). They catalyze key steps of pathways for the biosynthesis of not only physiologically essential but also industrially useful metabolites, such as amino acids, nucleotides, and fatty acids (Mitsuhashi et al., 2004). A role for carbonic anhydrase in the intracellular pH regulation has also been demonstrated in some bacteria (Marcus et al., 2005).

*Brucella* species contain two different β-CA, first identified in *B. suis* 1330, as thus named _Bs1330_CAI and _Bs1330_CAII. Both CAs contain the amino acid residues involved in binding of the Zn ion (typical of the β family of CAs), as well as those involved in the catalytic site. Their activity has been verified *in vitro*, and it is slightly higher in _Bs1330_CAII than _Bs1330_CAI (Joseph et al., 2010; Joseph et al. 2011). Pérez-Etayo et al. (2018) compared CAI and CAII activity (activity defined empirically as that allowing growth in a normal atmosphere, the same definition used throughout this study) in several strains of *B. suis*, *B. abortus* and *B. ovis*, and determined that CAII is not functional in CO_2_-dependent *B. abortus* and *B. ovis,* thus establishing a correlation between CA activity and CO_2_ dependence. They also observed that CAI is active in *B. suis* 1330 or 513, but not in *B. abortus* 2308W, 292 and 544. Moreover, although an active CAI alone is enough to support CO_2_-independent growth of *B. suis* in rich media, it is not able to do it in minimal media, or to support CO_2_-independent growth of *B. abortus* at all. A similar result was also obtained by Varesio et al (2019) that identified BcaA_BOV_ (CAII) as the enzyme responsible for the growth of *B. ovis* in a standard, unsupplemented atmosphere (0.04% CO2), in this case, by whole genome sequencing of CO_2_-independent mutants. Interestingly, they also reported that a CO2 downshift *B. ovis* initiates a gene expression program that resembles the stringent response and results in transcriptional activation of its type IV secretion system. This shift is absent in *B. ovis* strains carrying a functional copy of carbonic anhydrase.

The classical biotyping mentioned above, despite its limitations and the emergence of new molecular approaches to identify and classify *Brucella* at different taxonomic levels, is still extensively used by reference laboratories, often side by side with the new molecular methods (Garin-Bastuji et al., 2014). However, although there is a known link between phenotype and its genetic cause in some traits like urease activity or erythritol sensitivity (Sangari et al., 2007; Sangari et al., 1994), there is still a gap between the information provided by the molecular methods and the phenotype of *Brucella* isolates. With the availability of more genome sequences, it should be possible to reduce this gap by comparing the phenotypic characteristics of *Brucella* strains with their genome content. Comparative genomics of whole-genome sequences is especially interesting in bacterial pathogenesis studies (Hu et al., 2011). Pathogenomics can be considered as a particular case of comparative genomics, and it has been extensively used for the identification of putative virulence factors in bacteria, by comparing virulent and avirulent isolates (Pallen and Wren, 2007), although in principle could be applied to the elucidation of any phenotypic trait. The genus *Brucella* is a very homogeneous one, with over 90% identity on the basis of DNA-DNA hybridization assays within the classical species, and this results in relatively minor genetic variation between species that sometimes result in striking differences. As an example, only 253 single nucleotide polymorphisms (SNPs) separate *B. canis* from its nearest *B. suis* neighbour (Foster et al., 2009), but their host specificity differs widely; while *B. canis* is almost entirely restricted to the *Canidae* family, *B. suis* has a wide host range that includes pigs, dogs, rodents, hares, horses, reindeer, musk oxen, wild carnivores and humans. Similarly, there are only 39 SNPs consistently different between the vaccine strain *B. abortus* S19 and strains *B. abortus* 9-941 and 2308, two well-known virulent isolates (Crasta et al., 2008). In the last years a large number of *Brucella* genomes representing all species and biovars have been sequenced, and all this wealth of information is already resulting in new molecular epidemiology and typing methods (O’Callaghan and Whatmore, 2011). We have tested the potential of pathogenomics to unveil phenotypic traits in *Brucella* by defining the pangenome / pseudogenes of a set of *Brucella* strains, and comparing it with the CO_2_ dependence of those strains. This process has allowed us to identify Carbonic Anhydrase (CA) II, as the enzyme responsible for growth of the bacteria at atmospheric CO_2_ concentrations, and extend the analysis to new species of *Brucella*. All the sequenced genomes of *Brucella* contain two β-CA genes, but only those that carry a defective β-CA II require supplemental CO_2_. Reversion of this phenotype happens *in vitro* at a low frequency and is accompanied by a compensatory mutation that results in a full-length β-CAII product. We have also tested the hypothesis that the presence of a truncated β-CAII would have a competitive advantage *in vivo*, as a way to explain why a mutation with such a low frequency could get fixed in some *Brucella* species and biovars. A competitive assay shows that one of such mutants is significantly enriched in a mouse model of infection when compared with its corresponding full-length β-CAII strain. This could explain why CO_2_-dependent strains are selected *in vivo*. The polymorphisms affecting β-CAII encoding genes allow the prediction of the CO_2_-dependence status of any given strain, thus having the potential to replace the classical assay to characterize *Brucella* isolates.

## Materials and Methods

### Bacterial strains and growth conditions

The bacterial strains and plasmids used in this work are listed in Table 1. *Brucella* strains were grown at 37°C for 48–96 hours in a 5% CO_2_ atmosphere in *Brucella* broth (BB) or agar (BA) medium (Pronadisa, Spain). Media were supplemented with 10% foetal bovine serum to grow *B. ovis*. All experiments with live *Brucella* were performed in a Biosafety Level 3 facility at the Department of Molecular Biology of the University of Cantabria, and animal infections with *Brucella* were conducted at the University of Cantabria animal facilities, also under BSL3 conditions.

**Table 1.** Strains and plasmids used in this study.

| Strains | Main characteristics | Reference |
| --- | --- | --- |
| ***Brucella*** |  |  |
| *B. abortus* 544 | Biotype 1, CO<sub>2</sub>-dependent | Alton et al., 1988 |
| *B. abortus* 86/8/59 | Biotype 2, CO<sub>2</sub>-dependent | Alton et al., 1988 |
| *B. abortus* Tulya | Biotype 3, CO<sub>2</sub>-dependent | Alton et al., 1988 |
| *B. abortus* 292 | Biotype 4, CO<sub>2</sub>-dependent | Alton et al., 1988 |
| *B. abortus* A-579 | Biotype 3, CO<sub>2</sub>-dependent | Alton et al., 1988 |
| *B. abortus* 2308W | Biotype 1, CO<sub>2</sub>-independent | Suárez-Esquivel et al. 2016 |
| *B. ovis* 63/290 | CO<sub>2</sub>-dependent | Alton et al., 1988 |
| *B. pinnipedialis* B2/94 | CO<sub>2</sub>-dependent | Foster et al., 1996 |
| *B. abortus* 2308<br>$\Delta_{Ba2308w}CAI$ mutant | CO<sub>2</sub>-independent | This study |
| *B. abortus* 292 mut 1 | CO<sub>2</sub>-independent | This study |

### Bioinformatic methods

Genomic and protein sequences of the different *Brucella* species were obtained from GenBank and the Broad Institute (https://www.broadinstitute.org/projects/brucella). To allow easy comparison between the genes and pseudogenes in the different *Brucella* species, we constructed the panproteome of a selected set of 10 strains with the most complete genome annotation at the time (Table S1). To construct this set we started with all the CDS annotated in the *B. suis* 1330 genome. Next we found the most probable functional counterparts for the n pseudogenes annotated in *B. suis* 1330. The pseudogene list was taken directly from the original annotation of the *B. suis* 1330 genome. Finally, we added those CDS in indels from the other genomes not present in *B. suis* 1330. We assigned a new gene name to every CDS in our set following the Bru1_xxxx and Bru2_xxxx nomenclature, depending on the location of the gene in the *B. suis* genome. CDS from indels were also renamed with a nomenclature, BRU1_iXXXX, the “i” indicating their origin from indels absent in *B. suis* 1330. The file pan_pep provided in the supplementary materials is a multifasta protein file containing the sequence of all the 3496 CDS present at least once in any of the used genomes, and constitutes the first version of the *Brucella* pan proteome. The genes and pseudogenes annotated in these genomes were tabulated and assigned to one of the different gene families present in those genomes. In this way we constructed a spreadsheet with the pseudogenes in each genome using a uniform nomenclature. The analysis of the CA sequences at both the DNA and protein levels was extended to a group of 35 *Brucella* genomes (Table S2 in the supplementary material).

A structural theoretical model of *Brucella* _Ba2308_CAII was generated by molecular threading using the protein homology and recognition engine Phyre2 (Kelley et al., 2015), taking the atomic coordinates of the best hitas template. The pdb model generated was visualized using the PyMOL Molecular Graphics System, version 1.3 (Schrödinger, LLC, Portland, OR, USA).

Primers used in this study (Table 2) were designed with Primer 3 (http://bioinfo.ut.ee/primer3-0.4.0/) and synthesized by Sigma-Aldrich.

**Table 2.**
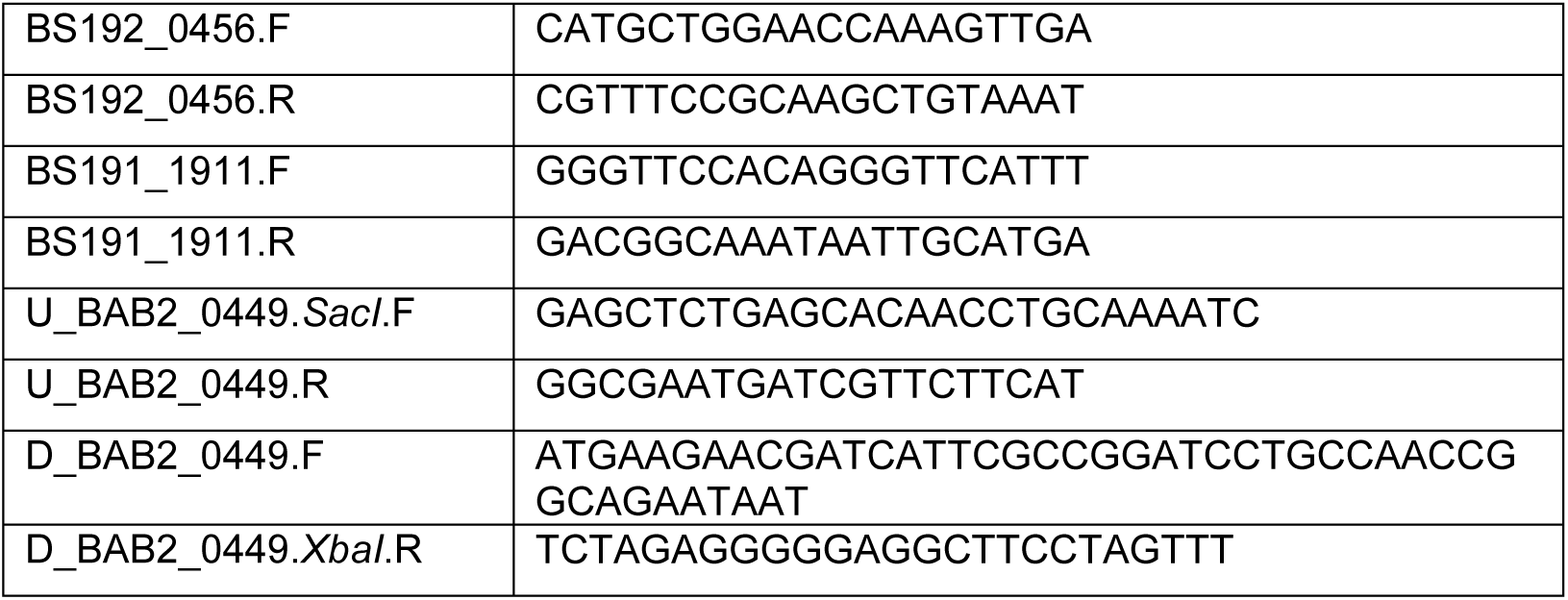
Oligonucleotides used in this work.

### Isolation of CO_2_-independent mutants in CO_2_-dependent *Brucella* strains

Different CO_2_-dependent *Brucella* strains from our collection were streaked onto BA plates and grown in a 5% CO_2_ atmosphere. Individual colonies were then re-streaked in duplicate plates, and incubated at 5% CO_2_, and ambient atmosphere to check for the correct CO_2_-dependence phenotype. They were grown as a lawn in fresh BA plates, and the growth was resuspended in PBS. The suspension was serially diluted and each dilution seeded in duplicate in BA plates. One dilution series was incubated at 5% CO_2_ to enumerate the number of bacteria in the inoculum, while the second was incubated at ambient atmosphere to select for CO_2_-independent colonies. The mutation rate was expressed as number of mutants per number of initial bacteria. Individual mutants were selected, and genomic DNA was obtained by using InstaGene matrix as described by the supplier (Bio-Rad Laboratories, United Kingdom). *CAI* and *CAII* complete sequences from the different strains were amplified by PCR with oligonucleotides BS192_0456.F/R and BS191_1911.F/R respectively, and sequenced to determine if there was any change compared to the corresponding parental sequence.

### Infection and intracellular viability assay of *B. abortus* in J774 cells

J774.A1 macrophage-like cells (ATCC, TIB-67) were cultured in RPMI medium with 2 mM L-glutamine, and 10% FBS at 37°C in 5% CO_2_ and 100% humidity. Confluent monolayers were trypsinized and 2×10^5^ cells/well were incubated for 24 h before infection in 24-well tissue culture plates. Macrophages were infected with *Brucella* strains in triplicate wells at a MOI of 50. After infection for 30 minutes, the wells were washed five times with sterile phosphate-buffered saline (PBS) and further incubated for 30 minutes in RPMI with 2 mM L-glutamine, 10% FCS and 50 µg gentamicin ml^−1^ to kill extracellular bacteria. That was taken as time 0 post-infection, and the medium was changed to contain 10 µg gentamicin ml^−1^. The number of intracellular viable *B. abortus* was determined at different time points by washing three times with PBS and lysing infected cells with 0.1% Triton X-100 in H_2_O and plating a series of 1:10 dilutions on BA plates for colony-forming unit (CFU) determination.

### Competitive infection assays

The following protocol was approved by the Cantabria University Institutional Laboratory Animal Care and Use Committee and was carried out in accordance with the Declaration of Helsinki and the European Communities Council Directive (86/609/EEC). Comparison of fitness between CO_2_-dependent and isogenic CO_2_-independent strains was done through a competitive infection assay in order to minimize animal-to-animal variation. BALB/c mice (CRIFA, Spain) were injected with 1:1 mixtures of *B. abortus* 292 (CO_2_ dependent, wild type) and *B. abortus* 292mut1 (a spontaneous *_Ba292_CAII* CO_2_-independent mutant). Two hundred microliters of a suspension containing approximately 10^8^ bacteria were administered intraperitoneally to a group (*n* = 6) of 6- to 8-week-old female BALB/c mice. Mice were sacrificed 8 weeks after infection, and the liver and spleen were removed aseptically and homogenized with 5 ml of BB containing 20% glycerol. Samples were serially diluted and plated in quadruplicate on BA plates. Half of the plates were incubated with 5% CO_2_, and the other half at ambient atmosphere. Additionally, colonies grown at 5% CO_2_ were replica-plated and incubated at both CO_2_ concentrations, to measure the ratio of CO_2_-dependent and CO_2_-independent colonies in two independent ways. For *in vitro* CI assays, BA plates were seeded forming a lawn with the infection mix, and incubated at 37°C with 5% CO_2_ for eight weeks, with repeated subculture in fresh BA plates every 4-5 days in the same conditions. The ratio of CO_2_-dependent and CO_2_-independent colonies was determined with the same protocol as the *in vivo* CI. The competitive index (CI) was calculated as the ratio of mutant to wild-type bacteria recovered at the end of the experiment divided by the ratio of mutant to wild-type bacteria in the inoculum, and the differences between groups were analyzed by Student’s two-tailed t test with significance set at P<0.05.

## Results

### Identification of the gene responsible for the CO_2_-dependence in *B. abortus*

The first evidence of the involvement of Carbonic Anhydrase in the CO_2_ dependence phenotype came from the analysis of pseudogenes in the ten fully annotated *Brucella* genomes (Table S1). After tabulation of the pseudogenes, their presence along with the different species was compared with the target phenotype, in this particular case we interrogated the spreadsheet n_pseudos.xls (Supplementary material) to find out which genes are pseudogenes only in those strains in our list that are CO_2_ dependent, *B. abortus* 9-941 and *B. ovis*. Three genes met this criterium, namely Bru1_1050 which encodes for a multidrug resistance efflux pump, Bru1_1827 which encodes for carbonic anhydrase II and Bru2_1236, encoding for an Adenosylmethionine-8-amino-7-oxononanoate aminotransferase.

Given the requirement of CA for growth of other microorganisms at ambient CO_2_ concentrations, and to check if Bru1_1827 could be responsible for the CO_2_-dependence phenotype, we retrieved and aligned the DNA and corresponding amino acid sequences obtained from a set of 35 *Brucella* strains with a known requirement for CO_2_ (Wattam *et al*, 2014), (Table S2). Sequences were clustered with VSEARCH (Rognes et al., 2016), resulting in 10 unique sequences that were aligned with ClustalW (Larkin et al., 2007). The CO_2_-independent isolates code for full-length identical proteins except for the *B. abortus* 2308 and 2308A strains that have an extra amino acid, Ala113. On the contrary, the CO_2_-dependent isolates contain different frameshifts or single point mutations, that result in truncated or altered proteins (Figure 1).

**Figure 1.**
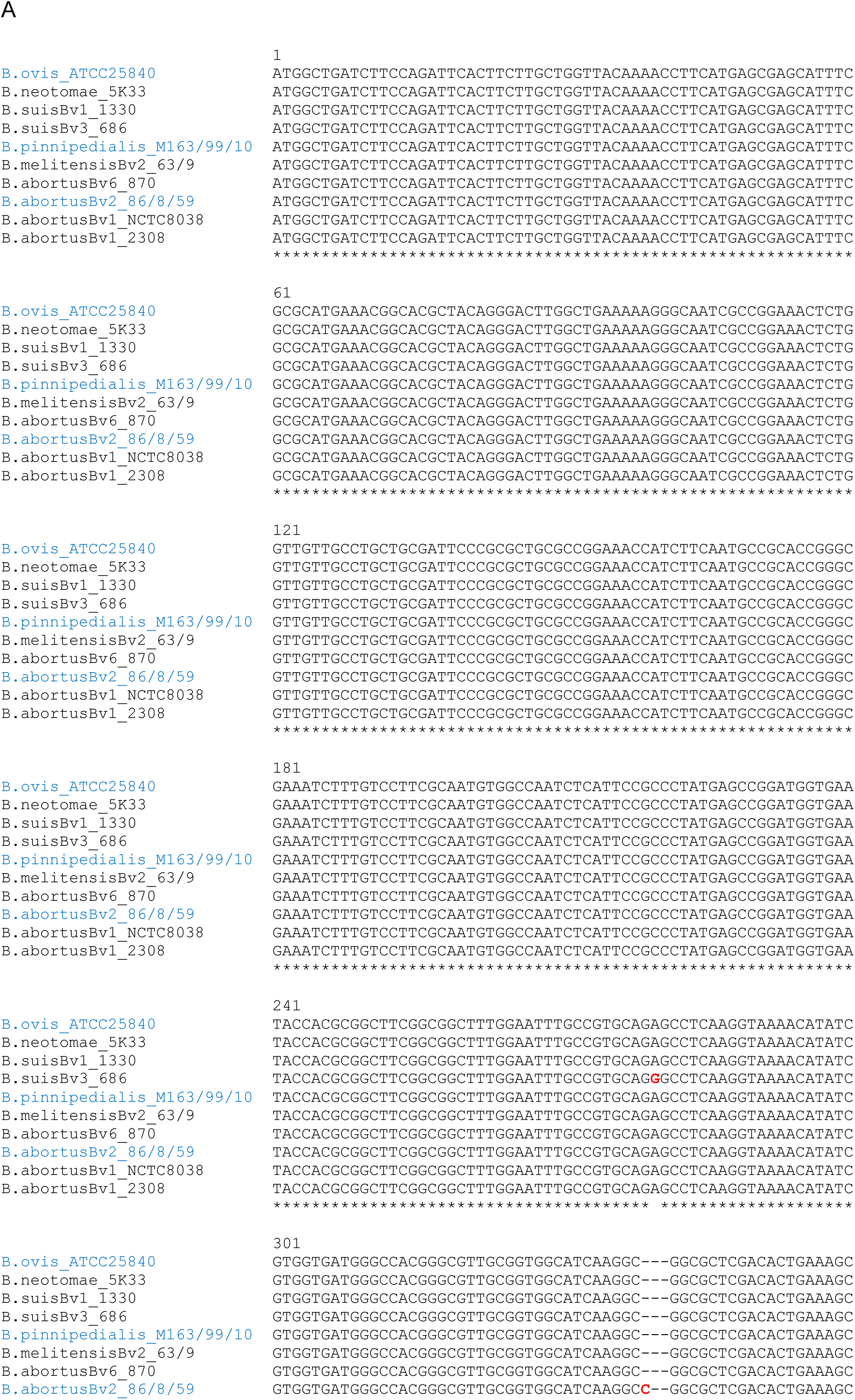
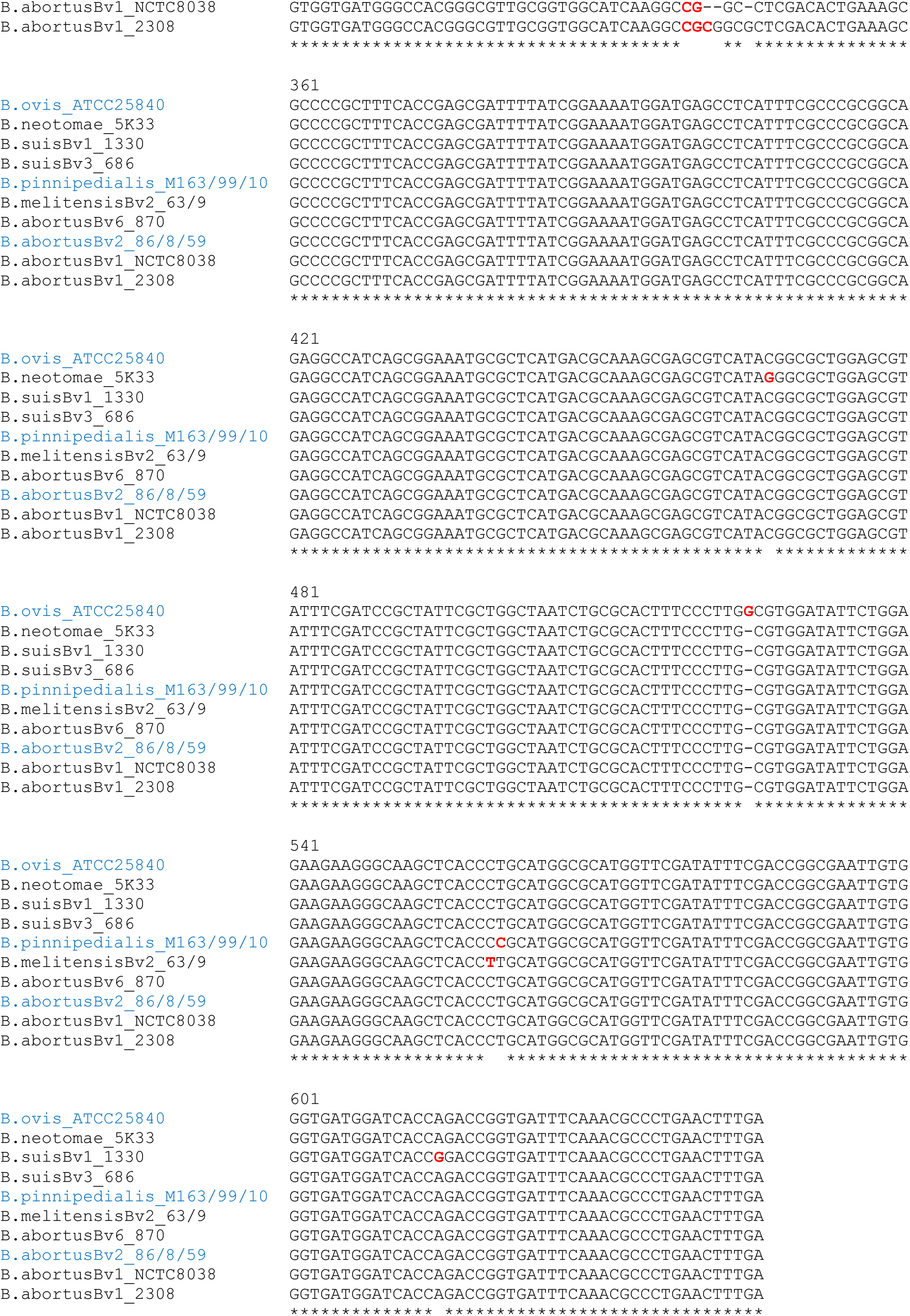
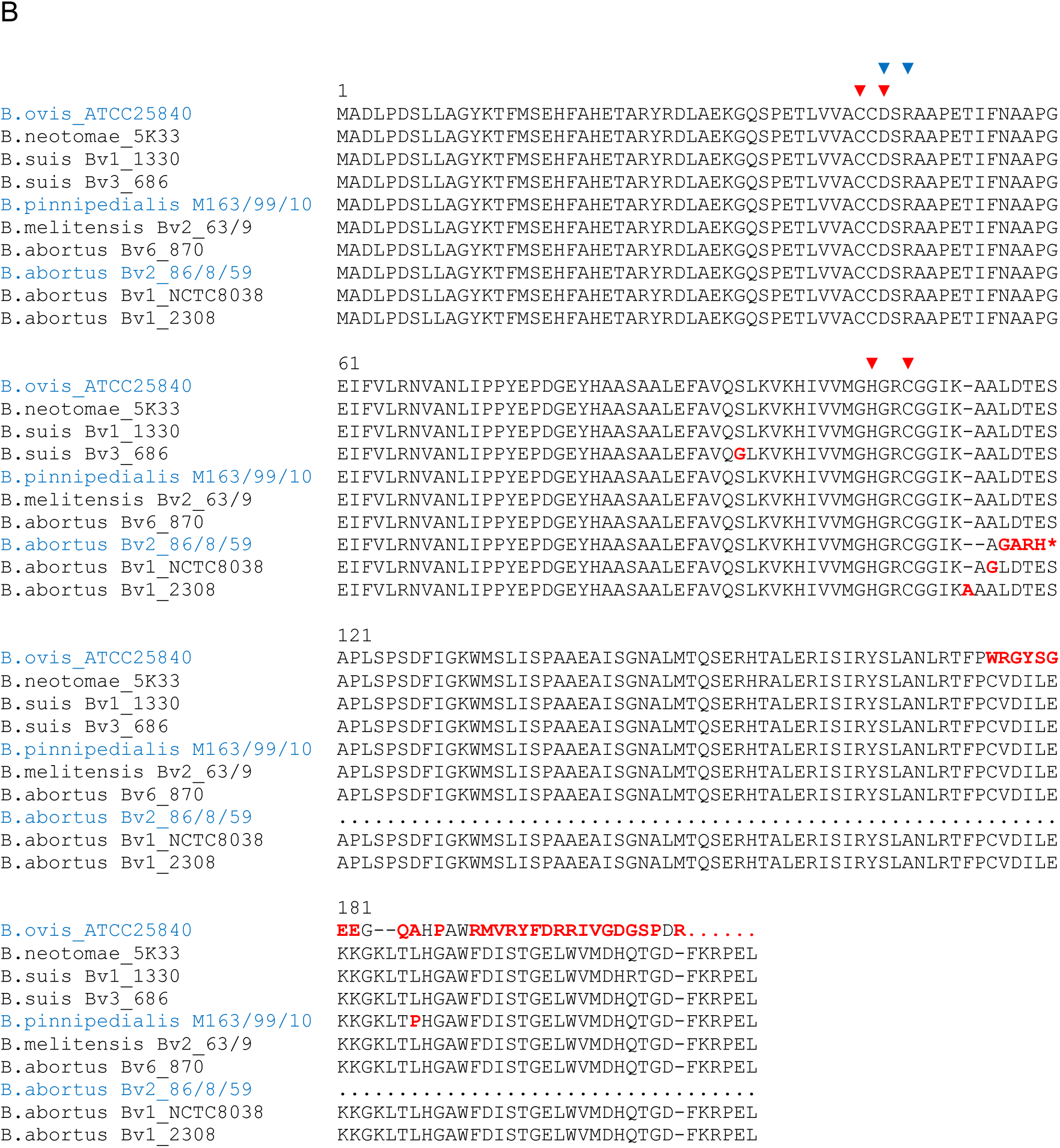
Alignment of sequences of Carbonic Anhydrase II from representative *Brucella* isolates. The genomes shown here are the representative species for each of the clusters of identical sequences obtained from the selected 35 *Brucella* strains. Those clusters formed by species that are CO_2_-dependent are shown in blue a) Partial DNA sequences, with nucleotides that differ from the consensus wild type sequence highlighted in red, and b) protein sequences, with amino acids that differ from the consensus wild type sequence highlighted in red. Red triangles indicate the four zinc-binding residues, Cys42, Asp44, His98, and Cys101 and blue triangles the catalytic dyad Asp44–Arg46

A group of 3 *B. abortus* strains (86/8/59, 9-941, and 292) shows an extra “C” at position 337 in the *CAII* gene when compared with the wild type allele, leading to a frameshift that causes a premature stop, truncating half of the protein. *B. ovis* ATCC 25840 shows an extra “G” at position 523, that similarly leads to a frameshift that alters the last third of the protein at the C-terminus. Finally, 3 *B. pinnipedialis* strains (M163/99/10, M292/94/1, and B2/94) contain a SNP, 557T>C, that causes a non-conservative amino acid substitution, Leu186Pro (Fig. 1B).

C337 also appears in CO_2_-independent *B. abortus* biovar 1 strains, like S19, 2308, or NTCC 8038, but in these cases there are additional mutations that recover the original ORF; two extra nucleotides in strain 2308, or one nucleotide deletions in *B. abortus* NCTC 8038 and S19. These changes do not affect the conserved amino acid residues typical of β-CAs involved in the catalytic cycle, that is, the four zinc-binding residues, Cys44, Asp46, Hys105, Cys108, and the catalytic dyad Asp46 and Arg48 (Fig. 1B). Some of the strains analyzed here (*B. suis* 1330, and *B. abortus* strains 2308W, 292 and 544) were also analyzed by Pérez-Etayo et al, (2018), and our results are in complete agreement.

There is one discrepancy involving *B. abortus* Tulya, a biovar 3 strain that according to the literature (Alton et al., 1988) should be CO_2_-dependent, but according to our analysis codes for a full-length CAII, thus being grouped with the CO_2_-independent isolates. To solve this apparent puzzle, we plated a sample of *B. abortus* Tulya from our laboratory stock and determined its CO_2_-dependence. Contrary to the original reference strain phenotype, and in agreement with our *in silico* analysis, this isolate was indeed CO_2_-independent. The complete *_BaTulya_CAII* was amplified by PCR from our strain and sequenced, confirming the published sequence. This strain originated from the collection kept in the Centro de Investigación y Tecnología Agroalimentaria of Aragón (CITA), Zaragoza, Spain, where it is also labelled as being CO_2_-independent, suggesting that this is not the result of a contamination or selection of a CO_2_-independent mutant in our hands.

As *Brucella* species code for two different carbonic anhydrases (Joseph et al., 2010), we repeated the analysis for the CAI-coding sequences (CDS). Although several isolates contain a polymorphism consisting of a 24 nt deletion between two 11 nt direct repeats, or different SNPs (Fig S1, Supplemental information), there was no obvious correlation between the presence of these polymorphisms and CO_2_-dependence. Pérez-Etayo et al (2018) demonstrated that CAI from *B. abortus* strains 2308W, 292 and 544 is inactive, while that from *B. suis* strains 1330 and 513 is active, although it can only mediate CO_2_-independence in complex media, and in a rather prototrophic host. Comparison of _Bsuis513_CAI and _Babortus2308W_CAI reveals a difference of only one amino acid, the valine at position 74 being replaced by a glycine.

### *Brucella* CO2-independent spontaneous mutants present a modified CAII sequence

Comparison of some of the *CAII* sequences of *B. abortus* biovar 1 CO_2_-independent strains like 2308, S19, or NCTC 8038 with those of the other *Brucella* CO_2_-dependent and independent isolates suggests that reversion of the CO_2_ requirement is coincidental with the introduction of compensatory mutations able to reverse the initial frameshift described above. CO_2_-independent mutants have been previously reported to appear at a low frequency (3 x 10^−10^) in cultures of CO_2_-dependent strains by subculturing *in vitro* in the absence of supplementary CO_2_ (Marr and Wilson, 1950). We measured the frequency of the reversion in six CO_2_-dependent strains from our laboratory collection, by growing duplicate cultures with or without CO_2_. We first checked the phenotype of all the strains by streaking them in a BA that was incubated without added CO_2_. All the strains but *B. abortus* Tulya, as reported above, failed to grow in these conditions, in agreement with the published phenotype. We then plated o/n cultures from the CO_2_-dependent strains to obtain colonies grown at ambient atmosphere, and calculated the frequency of revertants for those strains (Table 3). The *B. abortus* strains had a similar frequency to the one described by Mar and Wilson, 10^−8^ to 10^−10^, but *B. ovis* and *B. pinnipedialis* had a higher frequency of reversion, 10^−6^. In an exploratory effort to identify a possible cause for these differences in mutation rates, we analyzed the presence and identity between strains of the most obvious proteins that could be involved in this phenotype, like DNA polymerases, MutT, MutS, MutD, etc. Blastp analysis showed that, in all the cases, the protein was not only present in all strains, but had a 100% identity, so we could not find any difference that could explain our results. Maybe the analysis of the frequency of reversion in more CO_2_-dependent strains will reveal if this is a species, biovars or even isolate phenotype. We selected a few revertants from each strain, and amplified by PCR the CAI and CAII coding regions. The amplicons were then sequenced to determine if any compensatory mutation had appear in those loci. In all cases we found compensatory mutations in the same region, around nucleotides 333-343. All mutations in this hot spot resulted in full length CAII proteins (Figure 2), or in the case of *B. pinnipedialis*, a C to T change that reverts the Leu to Pro substitution. In this case, we also found the insertion of a nucleotide triplet (CGC or CCG) at the hot spot, that results in the addition of an extra amino acid, either Ala113 or Arg113. That is the same position wherethe extra codon in _Ba2308_CAII is located. Although some of the compensatory mutations appear several times, the most common situation was to find different mutations for the same sequence.

**Figure 2.**
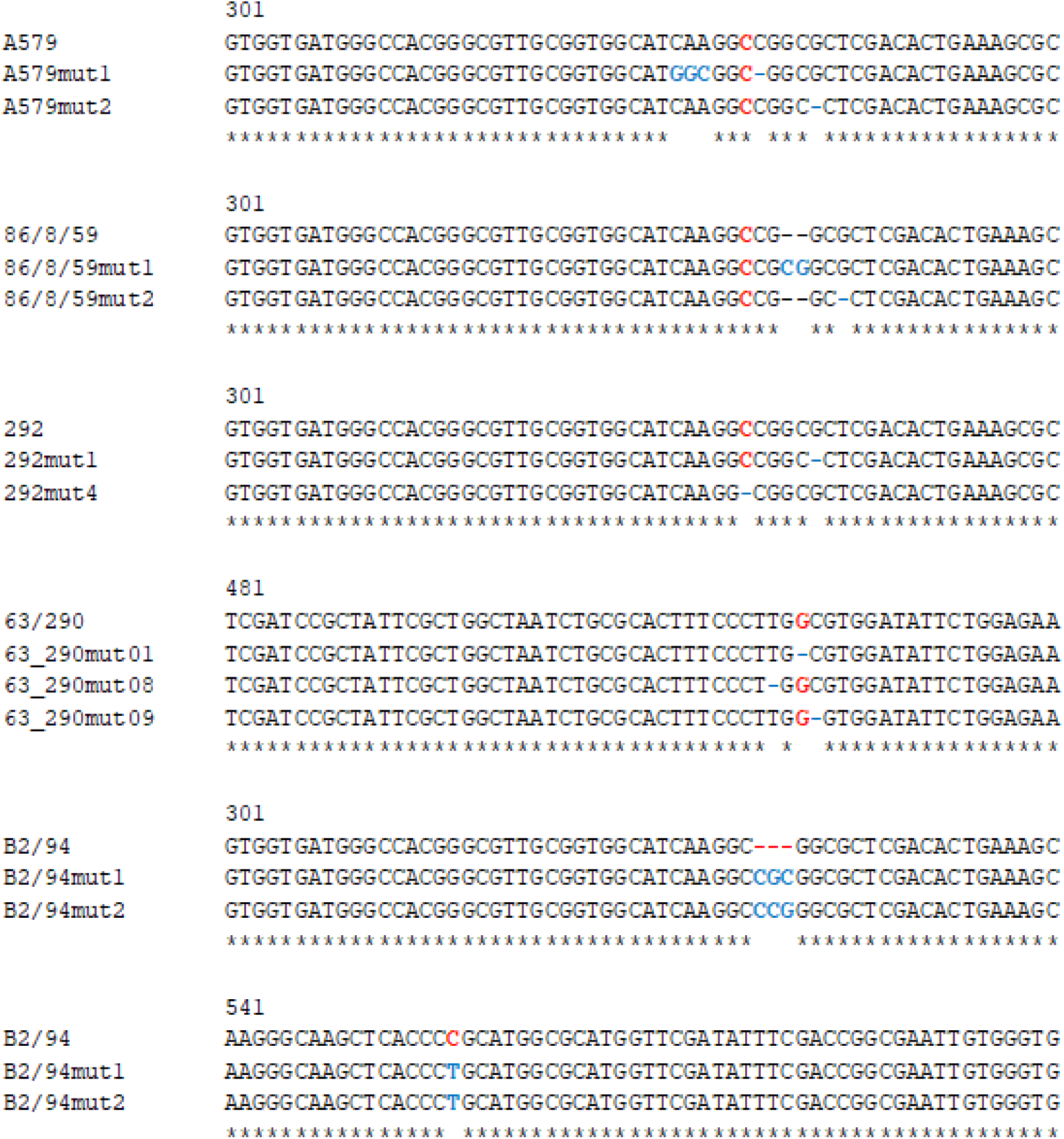
Nucleotide changes in CAII from selected CO_2_-independent mutants of different *Brucella* strains. Partial sequence of the regions where the original CO_2_-dependent strains had the mutations that caused the defective phenotype (shown in red), and the changes observed after selection and sequencing of different spontaneous CO_2_-independent mutants (shown in blue).

**Table 3.** Observed frequency of appearance of CO_2_-independent mutants in different *Brucella* strains.

| Strain | Frequency of mutants |
| --- | --- |
| *B. abortus* 292 | $2.72 \times 10^{-8}$ |
| *B. abortus* 544 | $5.95 \times 10^{-8}$ |
| *B. abortus* 86/8/59 | $1.14 \times 10^{-9}$ |
| *B. abortus* A-579 | $2.29 \times 10^{-10}$ |
| *B. ovis* 63/290 | $3.38 \times 10^{-6}$ |
| *B. pinnipedialis* B2/94 | $5.5 \times 10^{-6}$ |

As expected, reversion of the CO_2_-dependence phenotype did not produce any change in the coding sequence of CAI, reinforcing the hypothesis that CAII plays the main role in CO_2_-independence.

### Structural modelling of _Babortus2308_CAI and _Babortus2308_CAII

A single amino acid substitution, Val74 in _Bsuis513_CAI to Gly74 in _Babortus2308W_CAI, putatively renders the protein inactive, while the mutations in CAII in CO_2_-dependent *Brucella* isolates do not affect the region where the active center is located, at the N-ter part of the protein (Fig 1B). Moreover, a non-conservative Leu186Pro substitution, far from the active center, is enough as to induce CO_2_-dependence in the *B. pinnipedialis* strains analyzed. To better understand the effect of the observed mutations, a structural theoretical model of _Ba2308_CAI and _Ba2308_CAII was built with Phyre2. The modelled structures closely resembled those of other β-CAs that have been crystalized, displaying matches with a 100% confidence.

The closest structural homologue to _Ba2308_CAI is 1DDZ, a β-CA from the red alga *Porphyridium purpureum* (Mitsuhashi et al., 2000), with a 45% identity. Each 1DDZ monomer contains two internally repeated structures, each one homologous to _Ba2308_CAI. Overlapping of the modelled structures shows how the mutated residue Gly76 lies in close proximity to the coordinated zinc atom, and also to the dimer interface (Figure 3). In the equivalent position of Val76 in _Bsuis513_CAI, 1DDZ contains Ile173 or Ile427, both among the most hydrophobic of amino acids. These residues are stablishing hydrophobic contacts in the interface between the domains; Iso173 with Val441 and Phe442 (upper zoom image) and Iso427 with Phe168 and Tyr190 (lower zoom image). Identical (Phe71, Val90) and similar (Phe93) residues are located in the equivalent positions in *Brucella*_Ba2308_CAI. The presence of a Glycine in *Brucella*_Ba2308_CAI instead of an Isoleucine disrupts these hydrophobic interactions and could impair dimerization. Besides, this substitution could locally alter the folding of this region and affect the nearby residues that are coordinating the Zn atom. In both cases the structure, and consecuently the activity of the protein, would be affected. Indeed a Val to Gly substitution, located in the dimerization surface, was shown to interfere with dimerization of citrate synthase from *Thermoplasma acidophilum* (Kocabiyik and Erduran, 2000), reducing not only its catalytic activity (about 10-fold), but also decreasing its thermal and chemical stability.

**Figure 3.**
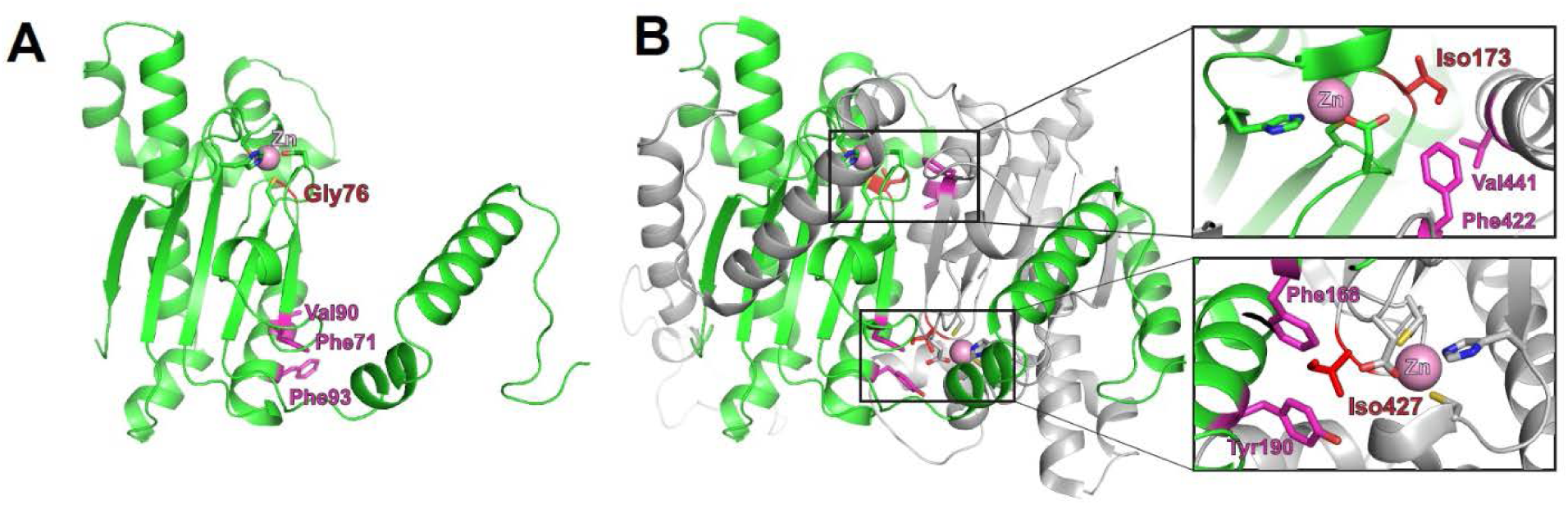
Structural model of *Brucella* _Ba2308_CAI. (A) Predicted structure of a monomer of Q2YL41, the CAI from *B. abortus* 2308, created using Phyre2 and the structure from the β-carbonic anhydrase from the red alga *Porphyridium purpureum* (1ddz) as template. Gly76 is depicted in red and the nearby zinc atom as a pink ball. (B) X-ray structure of the *Porphyridium purpureum* monomer, composed of two internally repeated structures. The N-terminal half (residues 1-308, equivalent to the sequence of monomeric _Ba2308_CAI) is colored in green and the C-terminal half (residues 309-564, equivalent to the second molecule of a putative dimer from _Ba2308_CAI) is colored in grey. In the equivalent position of Gly76 from _Ba2308_CAI in *Porphyridium purpureum* is located Iso173 (in the N-terminal half) or Iso427 (in the C-terminal half). These residues are stablishing hydrophobic contacts in the interface between the domains; Iso173 with Val441 and Phe442 (upper zoom image) and Iso427 with Phe168 and Tyr190 (lower zoom image). Identical (Phe71, Val90) and similar (Phe93) residues are located in the equivalent positions in *Brucella*_Ba2308_CAI (A). The presence of a glycine in *Brucella*_Ba2308_CAI instead of an isoleucine disrupts these hydrophobic interactions and could impair dimerization. Besides, this substitution could alter locally the folding of this region and affect the nearby residues that are coordinating the Zn atom.

The model structure obtained for _Ba2308_CAII is shown in Figure 4, along with the dimer structure of the best hit obtained, 5SWC, showing a 29% of identity and 100% confidence. 5SWC is the β-carbonic anhydrase CcaA from *Synechocystis* sp. PCC 6803. As _Ba2308_CAII contains an extra codon, the residue highlighted in red, Leu187, is the equivalent residue to the Leu186Pro change that is present in the CO_2_-dependent *B. pinnipedialis* strains.

**Figure 4.**
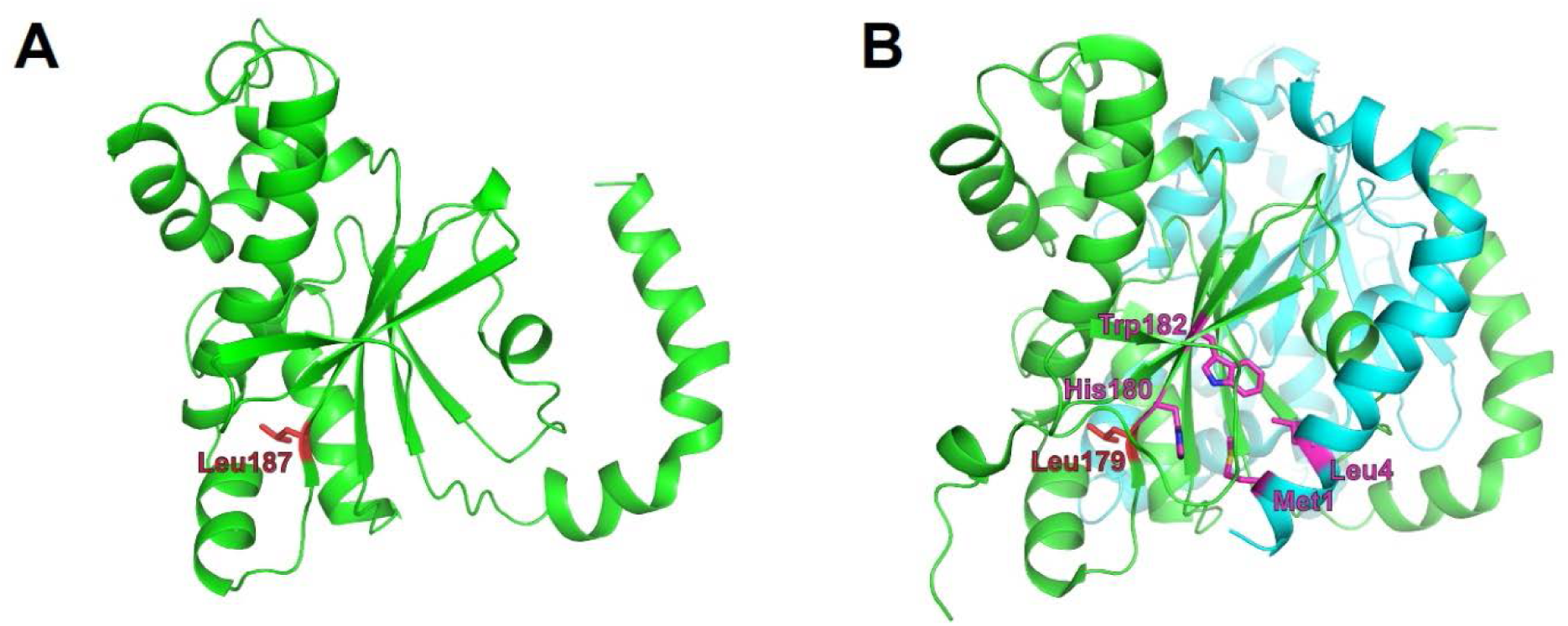
Structural theoretical model of *Brucella* _Ba2308_CAII. A) Model structure of a monomer of Q2YLK1, the CAII from *B. abortus* 2308, created using Phyre2 and structure 5SWC from *Synechocystis* sp as a template. Leu187 is depicted in red colour. (B) The X-ray structure of the *Synechocystis* sp CA dimer, showing an equivalent leucine residue in position 179. Adjacent residues His180 and Trp182 from the C-terminal β-sheet are in close contact with residues Met1 and Leu4 from the N-terminal H1-H2 helix, respectively, and are involved in monomer-monomer interactions. Zinc atoms are depicted as pink balls.

In this structure the protein crystalizes as a dimer, with the N-terminal arm composed of two alpha-helical segments (H1 and H2) that extend away from the rest of the molecule and make significant contacts with the last β-sheet with an adjacent monomer (in the case of _Ba2308_CAII His188 with Met1, and Trp191 with Leu4). This interaction between monomers has been determined as crucial for the establishment of the dimer (Cronk et al., 2001). In the case of *B. abortus* strains 86/8/59, 9-941, and 292, the premature stop would cause the complete loss of the C-ter end of the protein, including the last β-sheet, involved in the formation of the dimer. *B. ovis* ATCC 25840 shows also a completely altered C-terminus, and although the new amino acid sequence would remain folded as a β-sheet, it shows a completely different amino acid composition that would prevent the establishment of the right molecular interactions between the adjacent monomers. Regarding the last mutation observed in CO_2_-dependent strains, the SNP present in *B. pinnipedialis* strains M163/99/10, M292/94/1 and B2/94 causes a non-conservative amino acid substitution, Leu186Pro. The model predicts that this change will occur at the last β-sheet, in the area of interaction with the N-terminus of the adjacent monomer. Proline is an amino acid that confers an exceptional conformational rigidity, and as such is a known disruptor of both alpha helices and beta sheets. This being the case, this substitution is predicted to disrupt the dimerization of *Brucella* CAII.

### Competitive infection assays

Strain 2308 is not only a CO_2_-independent *Brucella* isolate, but also one of the most widely used virulent challenge strains, while S19, also a CO_2_-independent *Brucella* isolate is an attenuated vaccine strain. *In vitro* cell assays using J774 macrophages did not detect any difference in virulence between a CO_2_-dependent *B. abortus* 292 strain and its corresponding CO_2_-independent revertant (Figure 5). Additionally, we could not find any report in the literature that suggests that the CO_2_-dependence phenotype is related to virulence, and however there is one puzzling fact; despite the expected low frequency of a frameshift mutation, somehow this mutation is fixed in several species and biovars of *Brucella*. It is then reasonable to think of it as having a biological advantage in specific situations. Competitive index (CI) assays have been used to reveal subtle differences in fitness between two strains, and intra-animal experiments help to minimize inherent inter animal biological variation and also improve the identification of mutations or isolates with reduced or improved competitive fitness within the host (Falkow, 2004). As this could be the case with _Ba2308W_CAII, we performed a CI experiment using *B. abortus* 292 and one of its CO_2_-independent mutants, 292mut1. As a control we grew the same initial mixture in BA plates that were incubated at 37°C with 5% CO_2_, to know if any change in CI could be attributed to just the CO_2_ concentration, or there was some other factor that could be attributed to growth within an animal. Results are shown in Figure 6. During the course of the experiments in mice, there was a significant enrichment of the strain carrying the truncated form of _Ba2308W_CAII *B. abortus* 292, when compared with the CO_2_-independent revertant able to produce a complete active form of _Ba2308W_CAII. There was not a significant change in the ratio of both strains in liver or spleen, so the colony counts were combined in each mouse to show the ratio in that mice. At the same time, there was no significant enrichment / change in the ratio in cultures grown on plates. This suggests that inactivation of _Ba292_CAII has some fitness advantage *in vivo*, and could eventually result in the displacement of their corresponding CO_2_-independent counterpart. This hypothesis could explain why, despite the low frequency of mutation, CO_2_-dependent strains appear on primary isolation. As there are some other species and biotypes of *Brucella* that are CO_2_-dependent on primary isolation we could infer that the fitness advantage is also present in those species and biotypes.

**Figure 5.**
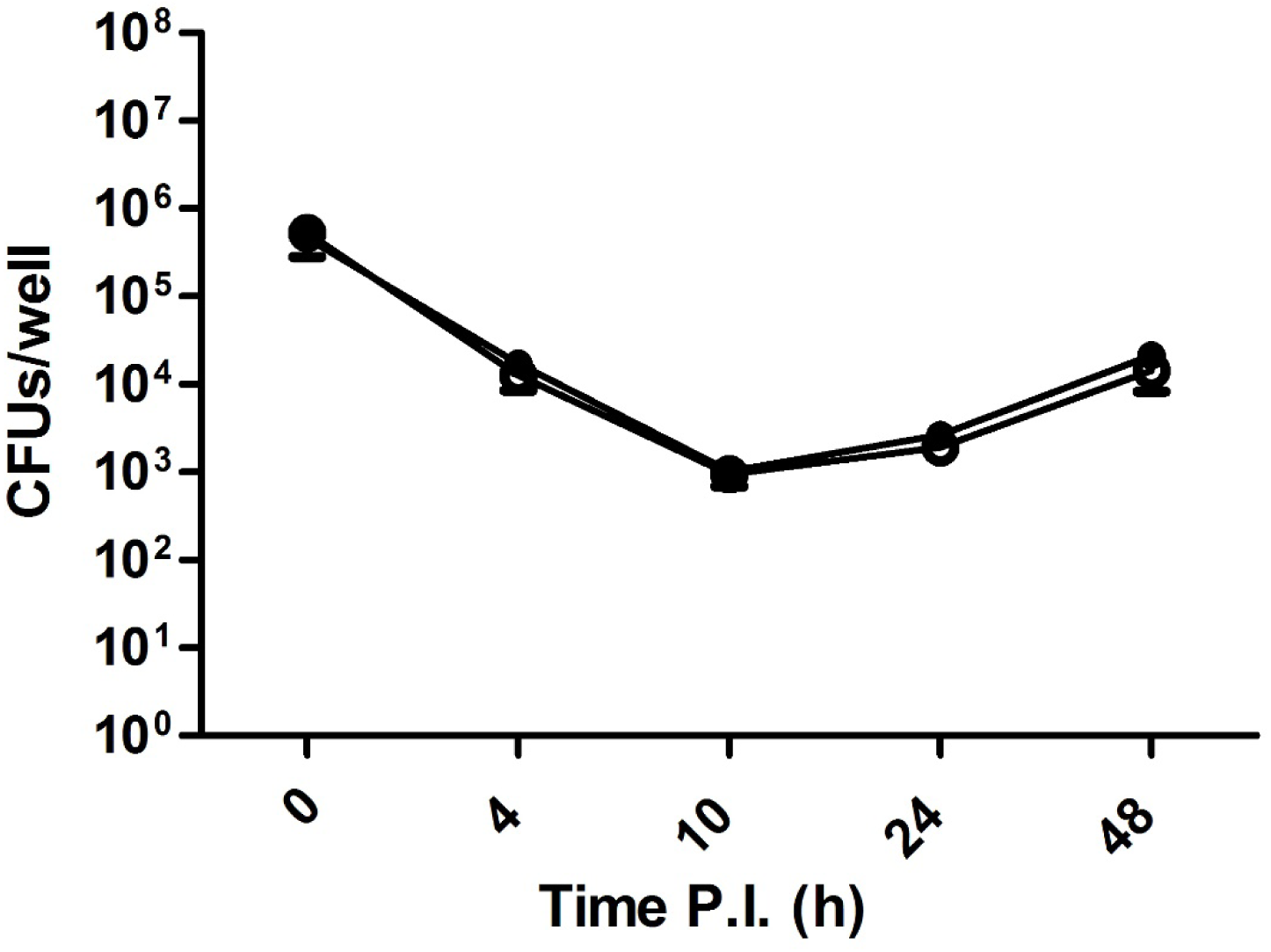
Infection and intracellular viability assay of *B. abortus* in J774 mouse macrophages *in vitro*. J774 macrophages were infected with either *B. abortus* 292 or the CO_2_-independent spontaneous mutant *B. abortus* 292mut1, at a MOI of 50. Samples from triplicate wells were obtained at 0, 4, 10, 24 and 48 h post infection, and enumerated by dilution and plating. ● *B. abortus* 292 ○ *B. abortus* 292mut1.

**Figure 6:**
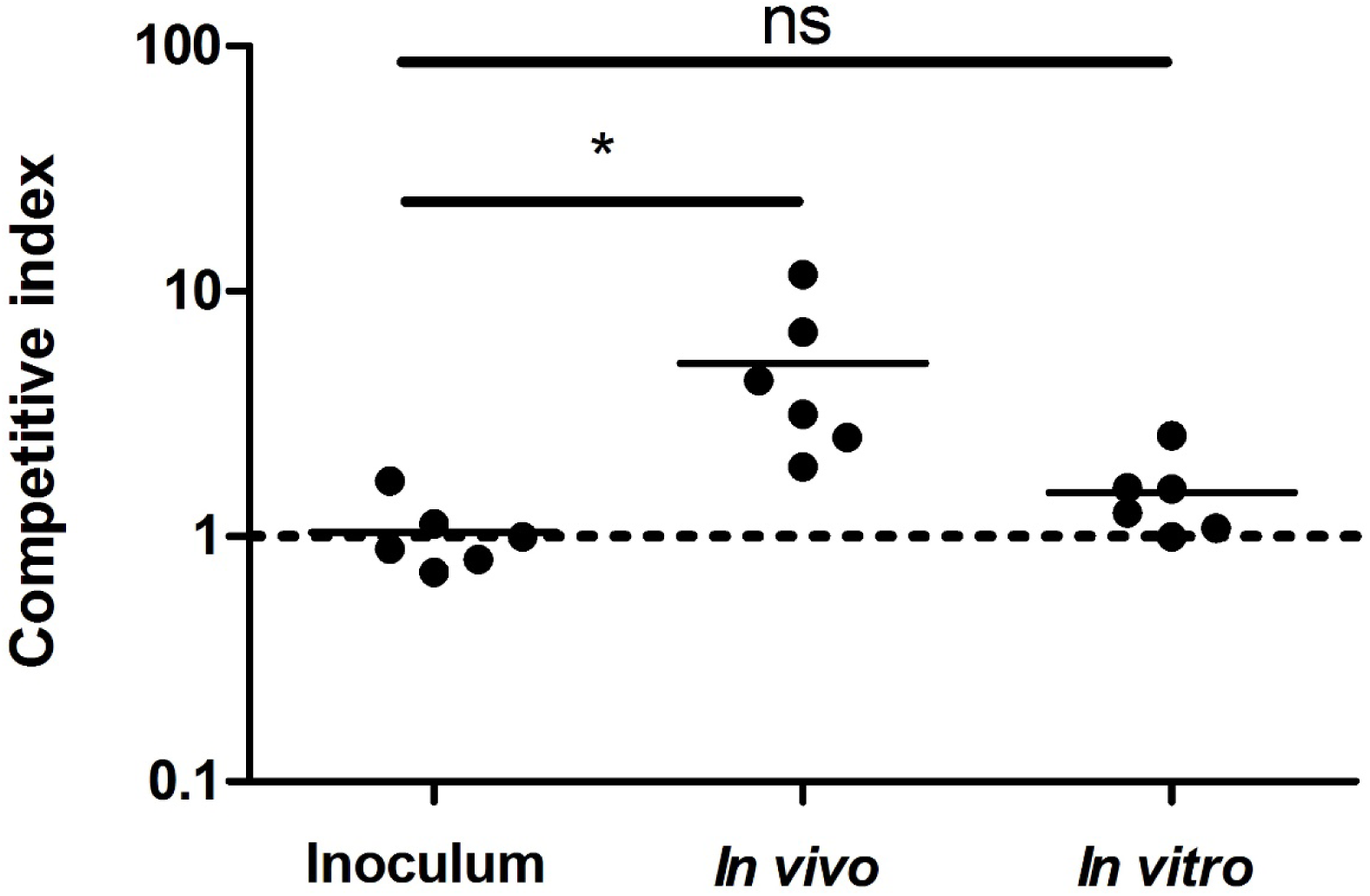
Competitive Index assay of a mixture of *B. abortus* 292, and its corresponding CO_2_-independent spontaneous mutant *B. abortus* 292 mut1. The competitive index was calculated by dividing the output ratio of mutant to wild-type bacteria by the input ratio of mutant to wild-type bacteria, in the two groups tested, regarding the original inoculum. Thus, for strains with the same fitness, the result should be 1. The differences between groups were analyzed by Student’s two-tailed t test with significance set at P<0.05 (*). ns, non significant.

## Discussion

Diagnosis of brucellosis is usually achived by serological detection in both animals and humans. This could be enough to warrant the initiation of response measures, like start of antibiotic therapy in humans, or immovilization or sacrifice of animals. However, isolation, identification and subtyping of brucellae is not only definitive proof of infection, but also allows epidemiological surveillance. Depending on the laboratory, this process is carried out by a combination of classical and modern molecular methods. The classical typing methods consist in the phenotypic characterization of the isolates, using biochemical and immunological tests (CO_2_ requirement, H_2_S production, urease activity, agglutination with monospecific A, R, and M sera, growth on media with thionin or basic fuchsin, or sensitivity to erythritol), and susceptibility to lytic *Brucella* phages (Alton et al., 1988). These methods require culture of the bacteria, are usually time-consuming and laborious, and they do not offer a good discriminatory power. Moreover, in the last years the field has experienced a revolution with the advent of new molecular methods, resulting in the description of new species, and a better understanding of the population structure of the genus *Brucella*. Thus the classical methods are being replaced or complemented by modern molecular methods. These methods range from PCR detection systems targeting different locus (like *ery*, *bcsp31*, or IS711), that allow species and even biovar differentiation (Mayer-Scholl et al., 2010; López-Goñi et al., 2011), to the multilocus sequence analysis (MLSA) that has been successfully used to describe the phylogenetic relationships of isolates, and the global population structure of the genus *Brucella* (Whatmore et al., 2016). More recently, with the advent of Whole Genome Sequencing (WGS), and especially with the drop in sequencing prices, WGS has been proposed to be the new routine typing method, particularly in groups with a high degree of similarity at the biochemical or serological levels (Chattaway et al. 2017), like *Brucellaceae* is. But these methods are still far from being routine in most brucellosis laboratories, particularly in developing countries, and the classical methods are still routinely used in reference laboratories. Although genomic information offers the potential to unveil most of the phenotypic traits in bacteria, there are still important attributes that are not evident in the genome sequence. Thus, there is a gap between the classical typing scheme and the molecular methods, and some features still can not be attributed to any specific genetic trait. In the case of *Brucella* it is particularly interesting the host specificity, that it is yet impossible to predict from the genome sequence. It is reasonable to think that as molecular typing improves we should advance in closing the gaps between classical and molecular typing, and we would be able to predict the full virulence and host specificity of a given isolate by analyzing the genome content. We have started to address this gap by looking at the genomic differences between *Brucella* isolates regarding one of the classical test for typing, as it is CO_2_ requirement.

*B. abortus* biovars 1, 2, 3, 4, and some isolates from biovar 9, as well as *B. ovis*, require an increased concentration of CO_2_ for growth, as do most strains of *B. pinnipedialis*, but only some of *B. ceti.* We selected 10 *Brucella* strains that have been sequenced and annotated, and which CO_2_ dependence status was known, to construct a *Brucella* pangenome based on the *B. suis* 1330 genome annotation. This resulted in a collection of 3496 CDS. We next compared the distribution of pseudogenes (as annotated in the databases) and absent genes with the CO_2_ dependence, resulting in only three candidate genes, Bru1_1050 which encodes for a multidrug resistance efflux pump, Bru1_1827 which encodes for carbonic anhydrase II and Bru2_1236, encoding for an Adenosylmethionine-8-amino-7-oxononanoate aminotransferase. The most obvious candidate was CAII, as it has been shown to be required to grow under ambient air in a number of microorganisms. To confirm our initial result, we extracted and aligned the DNA and amino acid sequences of CA II from an extended set of sequenced strains with a known CO_2_ phenotype. Those strains that are able to grow in atmospheric concentrations of CO_2_ carry a full length copy of the protein, while those that are not contain truncated or mutated versions of the proteins. *Brucella* species also carry a second carbonic anhydrase, CAI, but the polymorphisms found both at the DNA and protein levels do not allow to infer the CO_2_-dependence. This result is in agreement of that reported by Pérez-Etayo et al. (2018), and Varesio et al. (2019), and further extends the range of strains tested.

A direct application of this result would be the determination of the CO_2_-dependence status of any given strain by determining the sequence at the CAII locus. This is actually the case in *B. abortus* Tulya, where our analysis predicted that our stock should be CO_2_-independent, as it was the original stock from CITA. Laboratory determination of the phenotype confirmed the *in silico* result. This approach could be used to determine, or at least narrow down candidate genes for different phenotypes, obviously with monogenic traits being the easier to determine. We have found three different mutations that caused dependence of added CO_2_, two independent insertions (C337 and G523) that either cause a premature stop, or change completely the C-terminus of the protein, and a SNP that changes a leucine for a proline in the last β-sheet. All bacterial β-CAs crystallized so far are active as dimers or tetramers, and inactive as monomers, and all of them have the N-terminal α-helix arm that extends away from the rest of the molecule and makes significant contact with the last β-sheet of an adjacent monomer (Supuran, 2016). In all the cases observed in this work, the mutations do not affect the active site, but all of them potentially change the sequence and structure of the protein at the C-terminus so the most obvious hypothesis is that it is the modified structure of the proteins the cause for the loss of activity. Inactive *Brucella* CAII proteins either lack the last β-sheet completely, or have a very different sequence composition that disrupts this last β-sheet. The substitution of a leucine by a proline in the β-sheet is a particular example of this later case, as proline is known to be very disruptive amino acid for both α-helices and β-sheets structures. As these contacts seem to be important for dimerization, we can hypothesize that all the mutations found in CAII will have a strong impact in the dimerization or multimerization of CAII that will remain as a monomer, losing its activity (that we have defined as that allowing growth in a normal atmosphere). But there is a caveat in this reasoning. We, as well as others (Pérez-Etayo et al., 2018) have been unable to obtain a full-length mutant of CAII, despite being able to obtain a CAI (both data not shown). Moreovera transposon sequencing analysis shows that CAII is essential, at least for *B. abortus* 2308 (Sternon et al., 2018). This experiment was apparently carried out without added CO_2_, so the result is not unexpected. It would be interesting to know if, performed in the presence of 5-10% CO_2_, they would have observed insertions only in the C-ter of the protein, where the mutations in the natural CO_2_-dependent isolates accumulate. This means that the C-terminal part of the protein still carries out at least some of its functions as a monomer. We have not found any information regarding the activity of β-carbonic anhydrases as monomers, but in the α-carbonic anhydrase from *Thermovibrio ammonificans* the destabilization of the tetramer by reduction of the cysteines results in the dissociation of the tetrameric molecule into monomers with lower activity and reduced thermostability. It seems reasonable to think that this is the case also for *Brucella* CAII. CAII catalizes the fixation of CO_2_ with high efficiency when forming dimers, but the low efficiency of the carboxylation reaction when acting as a monomer would require the presence of higher amounts of CO_2_.

A similar situation could be taking place in the case of CAI. Modelling of the structure of _Babortus2308_CAI allows to hypothesize the role of the only residue of difference with _Bsuis513_CAI, that has to be responsible for the absence of activity in the first one. Its localization close to the Zn atom and to the dimer interface probably results in the destabilization of the dimer, lowering or abolishing its activity. However, it would be necessary to purify and characterized biochemically the monomers of both _Babortus2308_CAI and _Babortus2308_CAII to confirm our model.

These mutations can only be selected in high CO_2_ environments, like those present inside animals, where high CAII activity would be dispensable, as this atmosphere generates enough bicarbonate in solution as to fullfil the metabollic requirement of the bacteria (Nishimori 2009). We have determined the frequency of appearance of CO_2_-independent isolates, and although there is a huge variation between strains, it ranges from 10^−6^ to 10^−10^, as previously described. Despite its low frequency, somehow these mutations got selected in several species and biovars of *Brucella*, suggesting that they provide some biological advantage. To test this hypothesis, we performed a competitive assay both *in vitro* and *in vivo*. This assay resulted in a significant enrichment of the strain carrying an inactive carbonic anhydrase in animals, but not in cultured plates. Pérez-Etayo et at (2018) assayed the bacterial loads of *B. ovis* PA and *B. ovis* PA Tn7_Ba2308W_CAII in the spleens of BALB/c mice at 3 and 8 weeks post-infection, and found that there was no significant difference between a CO_2_-dependent and its corresponding CO_2_-independent strain at the level of multiplication in the mouse model. This apparent contradiction with our own results could be due to the different species used, or to the different experiment used to test this hypothesis. When trying to determine subtle differences in fitness between two given strains, a competitive assay has a higher discrimination power (Eekels et al., 2012; Shames et al, 2018), as any effect is amplified over time. Although the ultimate reason behind this competitive advantage is currently unknown, it would explain why some strains and biovars of *Brucella* are dependent of CO_2_ in primary isolation, despite the low frequency of mutation. It is also noteworthy that this phenotype is only observed in certain species and biovars, suggesting that the competitive advantage of the CAII mutants only applies to a subset of host/pathogen pairs. As CAII is essential, the mutant strains still would have to produce the protein, and thus the metabolic gain should be negligible for them. Another possibility would be that the dimer form of the enzyme is too active in a high CO_2_ environment, and causes a deleterious acidification in the bacteria. By evolving this sophisticated system that reversibly alters the dimerization state of the protein, *Brucella* is able to adjust to the different requirements encountered during its biological cycle.

## Supporting information

Pangenome annotation

## Acknowledgements

This work was supported by grants BFU2011-25658 from the Spanish Ministry of Science and Innovation, and by grant 55.JU07.64661 from the University of Cantabria to FJS. BAR was supported by a Scholarship received from DGAPA-UNAM. The authors want to acknowledge help from María J. Lucas and Elena Cabezón in the drawing and interpretation of crystallographic data.

