## Supplemental material for "Polymorphisms in *Brucella* Carbonic anhydrase II mediate CO_2_ dependence and fitness *in vivo*"

**Supplementary information**

**Excel File bru_pseudo.xlsx.** Sheet 1 (Pangenome) contains the complete list of genes found in the 10 genomes including their predicted function and functional categories. They have been named as described in the text. Sheet 2 (Pseudogenes by strain) contains the list of pseudogenes annotated in each one of the 10 genomes, using universal *Brucella* gene names as used in the Pangenome. Sheet 3 (Pseudogene list) contains the 726 genes which are pseudogenized at least in one of the 10 used genomes including annotation of the functional version of the gene. Sheet 4 is the phenotype calculator that allows to find combinations of pseudogenes by ordering the different columns.

**Table S1. *Brucella* genomes used to construct the pangenome and the pseudogene-phenotype predictor**

Accesion number	Pseudogenes

*Brucella abortus* bv. 1 str 9-941	NC_006932	NC_006933	169

*Brucella melitensis* bv. Abortus 2308	NC_007618	NC_007624	208

*Brucella abortus* S19	NC_010742	NC_010740	163

*Brucella melitensis* 16M	NC_003317	NC_003318	142

*Brucella melitensis* ATCC 23457	NC_012441.1	NC_012442.1	139

*Brucella suis* 1330	NC_004310	NC_004311	104

*Brucella suis* ATCC 23445	NC_010169.1	NC_010167.1	119

*Brucella ovis* ATCC 25840	NC_009505	NC_009504	205

*Brucella canis* ATCC 23365	NC_010103	NC_010104	100

*Brucella ceti* Cudo	(7 contigs, PRJNA33611)	144

**Table S2.** ***Brucella* genomes with identical Carbonic Anhydrase sequences.** Clustering of 35 *Brucella* strains (Genomes taken from Wattam *et al*, 2014) with a known requirement for CO_2_. Clusters of identical sequences were obtained with VSEARCH for a) *CAI*, and b) *CAII*. Highlighted in bold, the first strain of each cluster, that will be used as representative of the cluster for subsequent alignments (with the number of sequences belonging to that group in brackets).

A.

CO_2_ requirement	NCBI Bioproyect nr

**B.abortusBv1_2308 (7)**	PRJNA16203

B.abortusBv1_NCTC8038	PRJNA34743

B.abortusBv2_86/8/59	Yes	PRJNA243881
B.abortusBv4_292	Yes	PRJNA33027

B.abortusBv1_S19	PRJNA18999
B.abortusBv1_9-941	Yes	PRJNA9619
B.abortusBv1_2308A	PRJNA37723

**B.melitensisBv2_63/9 (7)**	PRJNA33577

B.pinnipedialis_B2/94	Yes	PRJNA33039
B.melitensisBv1_16MWGS	PRJNA34747
B.melitensisBv3_Ether	PRJNA33569
B.pinnipedialis_M292/94/1	Yes	PRJNA33563
B.neotomae_5K33	PRJNA33567
B.pinnipedialis_M163/99/10	Yes	PRJNA33037

**B.spF5/99 (4)**	PRJNA33767
B.ceti_B1/94	PRJNA33573
B.ceti_Cudo	PRJNA33611
B.ceti_M490/95/1	PRJNA33571

**B.abortusBv5_B3196 (3)**	PRJNA24387
B.abortusBv9_C68	PRJNA243877
B.abortusBv6_870	PRJNA244260

**B.melitensisBv1_16M (3)**	PRJNA180
B.ovis_ATCC25840	Yes	PRJNA12514
B.melitensisBv1_Rev.1	PRJNA33565

**B.suisBv1_1330 (3)**	PRJNA320
B.suisBv3_686	PRJNA33035
B. canis_RM6/66	PRJNA243891

**B.ceti_M13/05/1 (2)**	PRJNA33043
B.ceti_M644/93/1	PRJNA33041

**B.abortusBv3_Tulya (1)**	PRJNA33029
**B.microti_CCM4915 (1)**	PRJNA32233
**B.suisBv2_ATCC23445 (1)**	PRJNA20371
**B.suisBv4_40 (1)**	PRJNA34745

**B.suisBv5_513 (1)**	PRJNA33033
**B.sp_NVSL_07-0026 (1)**	PRJNA36511

B.

CO_2_ requirement	NCBI Bioproyect nr

**B.abortusBv6_870 (15)**	PRJNA244260

B.abortusBv9_C68	PRJNA243877

B.abortusBv3_Tulya	PRJNA33029

B.sp_NVSL07-0026	PRJNA36511

B.ceti_M13/05/1	PRJNA33043

B.ceti_M644/93/1	PRJNA33041

B.sp_F5/99	PRJNA33767

B.ceti_B1/94	PRJNA33573

B.ceti_M490/95/1	PRJNA33571

B.ceti_Cudo	PRJNA33611

B.canis_RM6/66	PRJNA24389

B.suisBv4_40	PRJNA34745

B.suisBv2_ATCC23445	PRJNA20371

B.suisBv5_513	PRJNA33033

B.microti_CCM4915	PRJNA32233

**B.melitensisBv2_63/9 (4)**	PRJNA33577

B.melitensisBv3_Ether	PRJNA33569

B.melitensisBv1_16M	PRJNA180

B.melitensisBv1_Rev.1	PRJNA33565

**B.abortusBv1_NCTC80 (4)**	PRJNA34743

B.melitensisBv1_16MWGS	PRJNA34747

B.abortusBv1_S19	PRJNA18999

B.abortusBv5_B3196	PRJNA24387

**B.abortusBv2_86/8/59 (3)**	Yes	PRJNA243881

B.abortusBv1_9-941	Yes	PRJNA9619

B.abortusBv4_292	Yes	PRJNA33027

**B.pinnipedialis_M163/99/10 (3)**	Yes	PRJNA33037

B.pinnipedialis_M292/94/1	Yes	PRJNA33563

B.pinnipedialis_B2/94	Yes	PRJNA33039

**B.abortusBv1_2308 (2)**	PRJNA16203

B.abortusBv1_2308A	PRJNA37723

**B.neotomae_5K33 (1)**	PRJNA33567

**B.ovis_ATCC_25840 (1)**	Yes	PRJNA12514

**B.suisBv1_1330 (1)**	PRJNA320

**B.suisBv3_686 (1)**	PRJNA33035

**Figure S1.** Alignment of the 13 identical clustered carbonic anhydrase I sequences from Table S1. A) DNA sequences. Different nucleotides are highlighted in red. The 11 nucleotide DRs where the deletion takes place are highlighted in blue. B). Protein sequences. Amino acids that are different are highlighted in red. Red triangles indicate the four zinc-binding residues*.*

A

```
                           1
B.abortusBv1_2308A         ATGCCCATGAAGAACGATCATTCGCCAGACCAGCGCACTTTATCGGAGCTTTTCGAGCAT
B.abortusBv5_B3196         ATGCCCATGAAGAACGATCATTCGCCAGACCAGCGCACTTTATCGGAGCTTTTCGAGCAT
B.spF5/99                  ATGCCCATGAAGAACGATCATTCGCCAGACCAGCGCACTTTATCGGAGCTTTTCGAGCAT
B.melitensisBv1_16M        ATGCCCATGAAGAACGATCATTCGCCAGACCAGCGCACTTTATCGGAGCTTTTCGAGCAT
B.melitensisBv2_63/9       ATGCCCATGAAGAACGATCATTCGCCAGACCAGCGCACTTTATCGGAGCTTTTCGAGCAT
B.suisBv1_1330             ATGCCCATGAAGAACGATCATTCGCCAGACCAGCGCACTTTATCGGAGCTTTTCGAGCAT
B.ceti_M13/05/1            ATGCCCATGAAGAACGATCATTCGCCAGACCAGCGCACTTTATCGGAGCTTTTCGAGCAT
B.abortusBv3_Tulya         ATGCCCATGAAGAACGATCATTCGCCAGACCAGCGCACTTTATCGGAGCTTTTCGAGCAT
B.suisBv5_513              ATGCCCATGAAGAACGATCATTCGCCAGACCAGCGCACTTTATCGGAGCTTTTCGAGCAT
B.sp_NVSL07-0026           ATGCCCATGAAGAACGATCATTCGCCAGACCAGCGCACTTTATCGGAGCTTTTCGAGCAT
B.suisBv2_ATCC23445        ATGCCCATGAAGAACGATCATTCGCCAGACCAGCGCACTTTATCGGAGCTTTTCGAGCAT
B.microti_CCM4915          ATGCCCATGAAGAACGATCATTCGCCAGACCAGCGCACTTTATCGGAGCTTTTCGAGCAT
B.suisBv4_40               ATGCCCATGAAGAACGATCATTCGCCAGACCAGCGCACTTTATCGGAGCTTTTCGAGCAT
                           ************************************************************

                           61
B.abortusBv1_2308A         AACCGTCAATGGGCAGCAGAAAAGCAGGAGAAAGACCCTGAATATTTCAGCCGCCTGTCA
B.abortusBv5_B3196         AACCGTCAATGGGCAGCAGAAAAGCAGGAGAAAGACCCTGAATATTTCAGCCGCCTGTCA
B.spF5/99                  AACCGTCAATGGGCAGCAGAAAAGCAGGAGAAAGACCCTGAATATTTCAGCCGCCTGTCA
B.melitensisBv1_16M        AACCGTCAATGGGCAGCAGAAAAGCAGGAGAAAGACCCTGAATATTTCAGCCGCCTGTCA
B.melitensisBv2_63/9       AACCGTCAATGGGCAGCAGAAAAGCAGGAGAAAGACCCTGAATATTTCAGCCGCCTGTCA
B.suisBv1_1330             AACCGTCAATGGGCAGCAGAAAAGCAGGAGAAAGACCCTGAATATTTCAGCCGCCTGTTA
B.ceti_M13/05/1            AACCGTCAATGGGCAGCAGAAAAGCAGGAGAAAGACCCTGAATATTTCAGCCGCCTGTCA
B.abortusBv3_Tulya         AACCGTCAATGGGCAGCAGAAAAGCAGGAGAAAGACCCTGAATATTTCAGCCGCCTGTCA
B.suisBv5_513              AACCGTCAATGGGCAGCAGAAAAGCAGGAGAAAGACCCTGAATATTTCAGCCGCCTGTCA
B.sp_NVSL_07-0026          AACCGTCAATGGGCAGCAGAAAAGCAGGAGAAAGACCCTGAATATTTCAGCCGCCTGTCA
B.suisBv2_ATCC23445        AACCGTCAATGGGCAGCAGAAAAGCAGGAGAAAGACCCTGAATATTTCAGCCGCCTGTCA
B.microti_CCM4915          AACCGTCAATGGGCAGCAGAAAAGCAGGAGAAAGACCCTGAATATTTCAGCCGCCTGTCA
B.suisBv4_40               AACCGTCAATGGGCAGCAGAAAAGCAGGAGAAAGACCCTGAATATTTCAGCCGCCTGTCA
                           ********************************************************** *

                           121
B.abortusBv1_2308A         TCGTCGCAGCGCCCGGAATTTCTATGGATCGGCTGTTCGGACAGCCGCGTTCCGGCCAAT
B.abortusBv5_B3196         TCGTCGCAGCGCCCGGAATTTCTATGGATCGGCTGTTCGGACAGCCGCGTTCCGGCCAAT
B.spF5/99                  TCGTCGCAGCGCCCGGAATTTCTATGGATCGGCTGTTCGGACAGCCGCGTTCCGGCCAAT
B.melitensisBv1_16M        TCGTCGCAGCGCCCGGAATTTCTATGGATCGGCTGTTCGGACAGCCGCGTTCCGGCCAAT
B.melitensisBv2_63/9       TCGTCGCAGCGCCCGGAATTTCTATGGATCGGCTGTTCGGACAGCCGCGTTCCGGCCAAT
B.suisBv1_1330             TCGTCGCAGCGCCCGGAATTTCTATGGATCGGCTGTTCGGACAGCCGCGTTCCGGCCAAT
B.ceti_M13/05/1            TCGTCGCAGCGCCCGGAATTTCTATGGATCGGCTGTTCGGACAGCCGCGTTCCGGCCAAT
B.abortusBv3_Tulya         TCGTCGCAGCGCCCGGAATTTCTATGGATCGGCTGTTCGGACAGCCGCGTTCCGGCCAAT
B.suisBv5_513              TCGTCGCAGCGCCCGGAATTTCTATGGATCGGCTGTTCGGACAGCCGCGTTCCGGCCAAT
B.sp_NVSL_07-0026          TCGTCGCAGCGCCCGGAATTTCTATGGATCGGCTGTTCGGACAGCCGCTTTCCGGCCAAT
B.suisBv2_ATCC23445        TCGTCGCAGCGCACGGAATTTCTATGGATCGGCTGTTCGGACAGCCGCGTTCCGGCCAAT
B.microti_CCM4915          TCGTCGCAGCGCCCGGAATTTCTATGGATCGGCTGTTCGGACAGCCGCGTTCCGGCCAAT
B.suisBv4_40               TCGTCGCAGCGCCCGGAATTTCTATGGATCGGCTGTTCGGACAGCCGCGTTCCGGCCAAT
                           ************ *********************************** ***********
                           181
B.abortusBv1_2308A         GTGGTGACGGGCCTTCAGCCGGGCGAAGTCTTCGTCCACCGTAATGGCGCCAATCTCGTC
B.abortusBv5_B3196         GTGGTGACGGGCCTTCAGCCGGGCGAAGTCTTCGTCCACCGTAATGGCGCCAATCTCGTC
B.spF5/99                  GTGGTGACGGGCCTTCAGCCGGGCGAAGTCTTCGTCCACCGTAATGTCGCCAATCTCATC
B.melitensisBv1_16M        GTGGTGACGGGCCTTCAGCCGGGCGAAGTCTTCGTCCACCGT------------------
B.melitensisBv2_63/9       GTGGTGACGGGCCTTCAGCCGGGCGAAGTCTTCGTCCACCGTAATGTCGCCAATCTCGTC
B.suisBv1_1330             GTGGTGACGGGCCTTCAGCCGGGCGAAGTCTTCGTCCACCGTAATGTCGCCAATCTCGTC
B.ceti_M13/05/1            GTGGTGACGGGCCTTCAGCCGGGCGAAGTCTTCGTCCACCGTAATGTCGCCAATCTCGTC
B.abortusBv3_Tulya         GTGGTGACGGGCCTTCAGCCGGGCGAAGTCTTCGTCCACCGTAATGTCGCCAATCTCGTC
B.suisBv5_513              GTGGTGACGGGCCTTCAGCCGGGCGAAGTCTTCGTCCACCGTAATGTCGCCAATCTCGTC
B.sp_NVSL_07-0026          GTGGTGACGGGCCTTCAGCCGGGCGAAGTCTTCGTCCACCGTAATGTCGCCAATCTCGTC
B.suisBv2_ATCC23445        GTGGTGATGGGCCTTCAGCCGGGCGAAGTCTTCGTCCACCGTAATGTCGCCAATCTCGTC
B.microti_CCM4915          GTGGTGACGGGCCTTCAGCCGGGCGAAGTCTTCGTCCACCGTAATGTCGCCAATCTCGTC
B.suisBv4_40               GTGGTGACAGGCCTTCAGCCGGGCGAAGTCTTCGTCCACCGTAATGTCGCCAATCTCGTC
                           *******  *********************************

                           241
B.abortusBv1_2308A         CACCGTGCCGATCTCAACCTGCTTTCCGTTCTGGAATTCGCCGTCGGGGTTCTGGAAATC
B.abortusBv5_B3196         CACCGTGCCGATCTCAACCTGCTTTCCGTTCTGGAATTCGCCGTCGGGGTTCTGGAAATC
B.spF5/99                  CACCGTGCCGATCTCAACCTGCTTTCCGTTCTGGAATTCGCCGTCGGGGTTCTGGAAATC
B.melitensisBv1_16M        ------GCCGATCTCAACCTGCTTTCCGTTCTGGAATTCGCCGTCGGGGTTCTGGAAATC
B.melitensisBv2_63/9       CACCGTGCCGATCTCAACCTGCTTTCCGTTCTGGAATTCGCCGTCGGGGTTCTGGAAATC
B.suisBv1_1330             CACCGTGCCGATCTCAACCTGCTTTCCGTTCTGGAATTCGCCGTCGGGGTTCTGGAAATC
B.ceti_M13/05/1            CACCGTGCCGATCTCAACCTGCTTTCCGTTCTGGAATTCGCCGTCGGGGTTCTGGAAATC
B.abortusBv3_Tulya         CACCGTGCCGATCTCAACCTGCTTTCCGTTCTGGAATTCGCCGTCGGGGTTCTGGAAATC
B.suisBv5_513              CACCGTGCCGATCTCAACCTGCTTTCCGTTCTGGAATTCGCCGTCGGGGTTCTGGAAATC
B.sp_NVSL_07-0026          CACCGTGCCGATCTCAACCTGCTTTCCGTTCTGGAATTCGCCGTCGGGGTTCTGGAAATC
B.suisBv2_ATCC23445        CACCGTGCCGATCTCAACCTGCTTTCCGTTCTGGAATTCGCCGTCGGGGTTCTGGAAATC
B.microti_CCM4915          CACCGTGCCGATCTCAACCTGCTTTCCGTTCTGGAATTCGCCGTCGGGGTTCTGGAAATC
B.suisBv4_40               CACCGTGCCGATCTCAACCTGCTGTCCGTGCTGGAATTCGCCGTCGGGGTTCTGGAAATC
                                 ***************** ***** ******************************


                           301
B.abortusBv1_2308A         AAGCATATCATCGTTTGCGGACATTATGGCTGCGGTGGGGTGCGCGCGGCAATGGATGGC
B.abortusBv5_B3196         AAGCATATCATCGTTTGCGGACATTATGGCTGCGGTGGGGTGCGCGCGGCAATGGATGGC
B.spF5/99                  AAGCATATCATCGTTTGCGGACATTATGGCTGCGGTGGGGTGCGCGCGGCAATGGATGGC
B.melitensisBv1_16M        AAGCATATCATCGTTTGCGGACATTATGGCTGCGGTGGGGTGCGCGCGGCAATGGATGGC
B.melitensisBv2_63/9       AAGCATATCATCGTTTGCGGACATTATGGCTGCGGTGGGGTGCGCGCGGCAATGGATGGC
B.suisBv1_1330             AAGCATATCATCGTTTGCGGACATTATGGCTGCGGTGGGGTGCGCGCGGCAATGGATGGC
B.ceti_M13/05/1            AAGCATATCATCGTTTGCGGACATTATGGCTGCGGTGCGGTGCGCGCGGCAATGGATGGC
B.abortusBv3_Tulya         AAGCATATCATCGTTTGCGGACATTATGGCTGCGGTGGGGTGCGCGCGGCAATGGATGGC
B.suisBv5_513              AAGCATATCATCGTTTGCGGACATTATGGCTGCGGTGGTGTGCGCGCGGCAATGGATGGC
B.sp_NVSL_07-0026          AAGCATATCATCGTTTGCGGACATTATGGCTGCGGTGGGGTGCGCGCGGCAATGGATGGC
B.suisBv2_ATCC23445        AAGCATATCATCGTTTGCGGACATTATGGCTGCGGTGGGGTGCGCGCGGCAATGGATGGC
B.microti_CCM4915          AAGCATATCATCGTTTGCGGACATTATGGCTGCGGCGGGGTGCGCGCGGCAATGGATGGC
B.suisBv4_40               AAGCATATCATCGTTTGCGGACATTATGGCTGCGGCGGGGTGCGCGCGGCAATGGATGGC
                           *********************************** *  *********************

                           361
B.abortusBv1_2308A         TATGGCCATGGCATCATCGACAATTGGCTGCAACCCATTCGCGATATTGCGCAGGCCAAT
B.abortusBv5_B3196         TATGGCCATGGCATCATCGACAATTGGCTGCAACCCATTCGCGATATTGCGCAGGCCAAT
B.spF5/99                  TATGGCCATGGCATCATCGACAATTGGCTGCAACCCATTCGCGATATTGCGCAGGCCAAT
B.melitensisBv1_16M        TATGGCCATGGCATCATCGACAATTGGCTGCAACCCATTCGCGATATTGCGCAGGCCAAT
B.melitensisBv2_63/9       TATGGCCATGGCATCATCGACAATTGGCTGCAACCCATTCGCGATATTGCGCAGGCCAAT
B.suisBv1_1330             TATGGCCATGGCATCATCGACAATTGGCTGCAACCCATTCGCGATATTGCGCAGGCCAAT
B.ceti_M13/05/1            TATGGCCATGGCATCATCGACAATTGGCTGCAACCCATTCGCGATATTGCGCAGGCCAAT
B.abortusBv3_Tulya         TATGGCCATGGCATCATCGACAATTGGCTGCAACCCATTCGCGATATTGCGCAGGCCAAT
B.suisBv5_513              TATGGCCATGGCATCATCGACAATTGGCTGCAACCCATTCGCGATATTGCGCAGGCCAAT
B.sp_NVSL_07-0026          TATGGCCATGGCATCATCGACAATTGGCTGCAACCCATTCGCGATATTGCGCAGGCCAAT
B.suisBv2_ATCC23445        TATGGCCATGGCATCATCGACAATTGGCTGCAACCCATTCGCGATATTGCGCAGGCCAAT
B.microti_CCM4915          TATGGCCATGGCATCATCGACAATTGGCTGCAACCCATTCGCGATATTGCGCAGGCCAAT
B.suisBv4_40               TATGGCCATGGCATCATCGACAATTGGCTGCAACCCATTCGCGATATTGCGCAGGCCAAT
                           ************************************************************

                           421
B.abortusBv1_2308A         CAGGCGGAACTGGACACCATAGAAAACACGCAGGACCGGCTGGACCGGCTTTGCGAATTG
B.abortusBv5_B3196         CAGGCGGAACTGGACACCATAGAAAACACGCAGGACCGGCTGGACCGGCTTTGCGAATTG
B.spF5/99                  CAGGCGGAACTGGACACCATAGAAAACACGCAGGACCGGCTGGACCGGCTTTGCGAATTG
B.melitensisBv1_16M        CAGGCGGAACTGGACACCATAGAAAACACGCAGGACCGGCTGGACCGGCTTTGCGAATTG
B.melitensisBv2_63/9       CAGGCGGAACTGGACACCATAGAAAACACGCAGGACCGGCTGGACCGGCTTTGCGAATTG
B.suisBv1_1330             CAGGCGGAACTGGACACCATAGAAAACACGCAGGACCGGCTGGACCGGCTTTGCGAATTG
B.ceti_M13/05/1            CAGGCGGAACTGGACACCATAGAAAACACGCAGGACCGGCTGGACCGGCTTTGCGAATTG
B.abortusBv3_Tulya         CAGGCGGAACTGGACACCATAGAAAACACGCAGGACCGGCTGGACCGGCTTTGCGAATTG
B.suisBv5_513              CAGGCGGAACTGGACACCATAGAAAACACGCAGGACCGGCTGGACCGGCTTTGCGAATTG
B.sp_NVSL_07-0026          CAGGCGGAACTGGACACCATAGAAAACACGCAGGACCGGCTGGACCGGCTTTGCGAATTG
B.suisBv2_ATCC23445        CAGGCGGAACTGGACACCATAGAAAACACGCAGGACCGGCTGGACCGGCTTTGCGAATTG
B.microti_CCM4915          CAGGCGGAACTGGACACCATAGAAAACACGCAGGACCGGCTGGACCGGCTTTGCGAATTG
B.suisBv4_40               CAGGCAGAACTGGACACCATAGAAAACACGCAGGACCGGCTGGACCGGCTTTGCGAATTG
                           ***** ******************************************************

                           481
B.abortusBv1_2308A         AGCGTTTCATCGCAGGTGGAAAGCCTGTCACGCACGCCGGTTCTGCAATCGGCCTGGAAG
B.abortusBv5_B3196         AGCGTTTCATCGCAGGTGGAAAGCCTGTCACGCACGCCGGTTCTGCAATCGGCCTGGAAG
B.spF5/99                  AGCGTTTCATCGCAGGTGGAAAGCCTGTCACGCACGCCGGTTCTGCAATCGGCCTGGAAG
B.melitensisBv1_16M        AGCGTTTCATCGCAGGTGGAAAGCCTGTCACGCACGCCGGTTCTGCAATCGGCCTGGAAG
B.melitensisBv2_63/9       AGCGTTTCATCGCAGGTGGAAAGCCTGTCACGCACGCCGGTTCTGCAATCGGCCTGGAAG
B.suisBv1_1330             AGCGTTTCATCGCAGGTGGAAAGCCTGTCACGCACGCCGGTTCTGCAATCGGCCTGGAAG
B.ceti_M13/05/1            AGCGTTTCATCGCAGGTGGAAAGCCTGTCACGCACGCCGGTTCTGCAATCGGCCTGGAAG
B.abortusBv3_Tulya         AGCGTTTCATCGCAGGTGGAAAGCCTGTCACGCACGCCGGTTCTGCAATCGGCCTGGAAG
B.suisBv5_513              AGCGTTTCATCGCAGGTGGAAAGCCTGTCACGCACGCCGGTTCTGCAATCGGCCTGGAAG
B.sp_NVSL_07-0026          AGCGTTTCATCGCAGGTGGAAAGCCTGTCACGCACGCCGGTTCTGCAATCGGCCTGGAAG
B.suisBv2_ATCC23445        GGCGTTTCATCGCAGGTGGAAAGCCTGTCACGCACGCCGGTTCTGCAATCGGCCTGGAAG
B.microti_CCM4915          AGCGTTTCATCGCAGGTGGAAAGCCTGTCACGCACGCCGGTTCTGCAATCGGCCTGGAAG
B.suisBv4_40               AGCGTTTCATCGCAGGTGGAAAGCCTGTCACGCACGCCGGTTCTGCAATCGGCCTGGAAG
                            ***********************************************************

                           541
B.abortusBv1_2308A         GACGGAAAGGATATCATCGTCCATGGCTGGATGTATAATCTGAAAGATGGGCTACTGCGC
B.abortusBv5_B3196         GACGGAAAGGATATCATCGTCTATGGCTGGATGTATAATCTGAAAGATGGGCTACTGCGC
B.spF5/99                  GACGGAAAGGATATCATCGTCCATGGCTGGATGTATAATCTGAAAGATGGGCTACTGCGC
B.melitensisBv1_16M        GACGGAAAGGATATCATCGTCCATGGCTGGATGTATAATCTGAAAGATGGGCTACTGCGC
B.melitensisBv2_63/9       GACGGAAAGGATATCATCGTCCATGGCTGGATGTATAATCTGAAAGATGGGCTACTGCGC
B.suisBv1_1330             GACGGAAAGGATATCATCGTCCATGGCTGGATGTATAATCTGAAAGATGGGCTACTGCGC
B.ceti_M13/05/1            GACGGAAAGGATATCATCGTCCATGGCTGGATGTATAATCTGAAAGATGGGCTACTGCGC
B.abortusBv3_Tulya         GACGGAAAGGATATCATCGTCCATGGCTGGATGTTTAATCTGAAAGATGGGCTACTGCGC
B.suisBv5_513              GACGGAAAGGATATCATCGTCCATGGCTGGATGTATAATCTGAAAGATGGGCTACTGCGC
B.sp_NVSL_07-0026          GACGGAAAGGATATCATCGTCCATGGCTGGATGTATAATCTGAAAGATGGGCTACTGCGC
B.suisBv2_ATCC23445        GACGGAAAGGATATCATCGTCCATGGCTGGATGTATAATCTGAAAGATGGGCTACTGCGC
B.microti_CCM4915          GACGGAAAGGATATCATCGTCCATGGCTGGATGTATAATCTGAAAGATGGGCTACTGCGC
B.suisBv4_40               GACGGAAAGGATATCATCGTCCATGGCTGGATGTATAATCTGAAAGACGGGCTGTTGCGC
                           ********************* ************ ************ *****  *****

                           601
B.abortusBv1_2308A         GATATCGGTTGCGACTGCACCCGCAATGCCTTGCAATTTGCCTGCCAACCGGCAGAA
B.abortusBv5_B3196         GATATCGGTTGCGACTGCACCCGCAATGCCTTGCAATTTGCCTGCCAACCGGCAGAA
B.spF5/99                  GATATCGGTTGCGACTGCACCCGCAATGCCTTGCAATTTGCCTGCCAACCGGCAGAA
B.melitensisBv1_16M        GATATCGGTTGCGACTGCACCCGCAATGCCTTGCAATTTGCCTGCCAACCGGCAGAA
B.melitensisBv2_63/9       GATATCGGTTGCGACTGCACCCGCAATGCCTTGCAATTTGCCTGCCAACCGGCAGAA
B.suisBv1_1330             GATATCGGTTGCGACTGCACCCGCAATGCCTTGCAATTTGCCTGCCAACCGGCAGAA
B.ceti_M13/05/1            GATATCGGTTGCGACTGCACCCGCAATGCCTTGCAATTTGCCTGCCAACCGGCAGAA
B.abortusBv3_Tulya         GATATCGGTTGCGACTGCACCCGCAATGCCTTGCAATTTGCCTGCCAACCGGCAGAA
B.suisBv5_513              GATATCGGTTGCGACTGCACCCGCAATGCCTTGCAATTTGCCTGCCAACCGGCAGAA
B.sp_NVSL_07-0026          GATATCGGTTGCGACTGCACCCGCAATGCCTTGCAATTTGCCTGCCAACCGGCAGAA
B.suisBv2_ATCC23445        GATATCGGTTGCGACTGCACCCGCAATGCCTTGCAATTTGCCTGCCAACCGGCAGAA
B.microti_CCM4915          GATATCGGTTGCGACTGCACCCGCAATGCCTTGCAATTTGCCTGCCAACCGGCAGAA
B.suisBv4_40               GATATTGGTTGCGACTGCACCCGCAATGCCCTGCAATTTGCCTGCCAACCGGCAGAA
                           ***** ************************ **************************
```

B

```
                           1                                                ▼ ▼
B.abortusBv1_2308A         MPMKNDHSPDQRTLSELFEHNRQWAAEKQEKDPEYFSRLSSSQRPEFLWIGCSDSRVPAN
B.abortusBv5_B3196         MPMKNDHSPDQRTLSELFEHNRQWAAEKQEKDPEYFSRLSSSQRPEFLWIGCSDSRVPAN
B.spF5/99                  MPMKNDHSPDQRTLSELFEHNRQWAAEKQEKDPEYFSRLSSSQRPEFLWIGCSDSRVPAN
B.melitensisBv1_16M        MPMKNDHSPDQRTLSELFEHNRQWAAEKQEKDPEYFSRLSSSQRPEFLWIGCSDSRVPAN
B.melitensisBv2_63/9       MPMKNDHSPDQRTLSELFEHNRQWAAEKQEKDPEYFSRLSSSQRPEFLWIGCSDSRVPAN
B.suisBv1_1330             MPMKNDHSPDQRTLSELFEHNRQWAAEKQEKDPEYFSRLLSSQRPEFLWIGCSDSRVPAN
B.ceti_M13/05/1            MPMKNDHSPDQRTLSELFEHNRQWAAEKQEKDPEYFSRLSSSQRPEFLWIGCSDSRVPAN
B.abortusBv3_Tulya         MPMKNDHSPDQRTLSELFEHNRQWAAEKQEKDPEYFSRLSSSQRPEFLWIGCSDSRVPAN
B.suisBv5_513              MPMKNDHSPDQRTLSELFEHNRQWAAEKQEKDPEYFSRLSSSQRPEFLWIGCSDSRVPAN
B.sp_NVSL_07-0026          MPMKNDHSPDQRTLSELFEHNRQWAAEKQEKDPEYFSRLSSSQRPEFLWIGCSDSRFPAN
B.suisBv2_ATCC23445        MPMKNDHSPDQRTLSELFEHNRQWAAEKQEKDPEYFSRLSSSQRTEFLWIGCSDSRVPAN
B.microti_CCM4915          MPMKNDHSPDQRTLSELFEHNRQWAAEKQEKDPEYFSRLSSSQRPEFLWIGCSDSRVPAN
B.suisBv4_40               MPMKNDHSPDQRTLSELFEHNRQWAAEKQEKDPEYFSRLSSSQRPEFLWIGCSDSRVPAN
                           *************************************** ****.***********.***

                           61                                           ▼  ▼
B.abortusBv1_2308A         VVTGLQPGEVFVHRNGANLVHRADLNLLSVLEFAVGVLEIKHIIVCGHYGCGGVRAAMDG
B.abortusBv5_B3196         VVTGLQPGEVFVHRNGANLVHRADLNLLSVLEFAVGVLEIKHIIVCGHYGCGGVRAAMDG
B.spF5/99                  VVTGLQPGEVFVHRNVANLIHRADLNLLSVLEFAVGVLEIKHIIVCGHYGCGGVRAAMDG
B.melitensisBv1_16M        VVTGLQPGEVFVHR--------ADLNLLSVLEFAVGVLEIKHIIVCGHYGCGGVRAAMDG
B.melitensisBv2_63/9       VVTGLQPGEVFVHRNVANLVHRADLNLLSVLEFAVGVLEIKHIIVCGHYGCGGVRAAMDG
B.suisBv1_1330             VVTGLQPGEVFVHRNVANLVHRADLNLLSVLEFAVGVLEIKHIIVCGHYGCGGVRAAMDG
B.ceti_M13/05/1            VVTGLQPGEVFVHRNVANLVHRADLNLLSVLEFAVGVLEIKHIIVCGHYGCGAVRAAMDG
B.abortusBv3_Tulya         VVTGLQPGEVFVHRNVANLVHRADLNLLSVLEFAVGVLEIKHIIVCGHYGCGGVRAAMDG
B.suisBv5_513              VVTGLQPGEVFVHRNVANLVHRADLNLLSVLEFAVGVLEIKHIIVCGHYGCGGVRAAMDG
B.sp_NVSL_07-0026          VVTGLQPGEVFVHRNVANLVHRADLNLLSVLEFAVGVLEIKHIIVCGHYGCGGVRAAMDG
B.suisBv2_ATCC23445        VVMGLQPGEVFVHRNVANLVHRADLNLLSVLEFAVGVLEIKHIIVCGHYGCGGVRAAMDG
B.microti_CCM4915          VVTGLQPGEVFVHRNVANLVHRADLNLLSVLEFAVGVLEIKHIIVCGHYGCGGVRAAMDG
B.suisBv4_40               VVTGLQPGEVFVHRNVANLVHRADLNLLSVLEFAVGVLEIKHIIVCGHYGCGGVRAAMDG
                           ** ***********        ******************************.*******

                           121
B.abortusBv1_2308A         YGHGIIDNWLQPIRDIAQANQAELDTIENTQDRLDRLCELSVSSQVESLSRTPVLQSAWK
B.abortusBv5_B3196         YGHGIIDNWLQPIRDIAQANQAELDTIENTQDRLDRLCELSVSSQVESLSRTPVLQSAWK
B.spF5/99                  YGHGIIDNWLQPIRDIAQANQAELDTIENTQDRLDRLCELSVSSQVESLSRTPVLQSAWK
B.melitensisBv1_16M        YGHGIIDNWLQPIRDIAQANQAELDTIENTQDRLDRLCELSVSSQVESLSRTPVLQSAWK
B.melitensisBv2_63/9       YGHGIIDNWLQPIRDIAQANQAELDTIENTQDRLDRLCELSVSSQVESLSRTPVLQSAWK
B.suisBv1_1330             YGHGIIDNWLQPIRDIAQANQAELDTIENTQDRLDRLCELSVSSQVESLSRTPVLQSAWK
B.ceti_M13/05/1            YGHGIIDNWLQPIRDIAQANQAELDTIENTQDRLDRLCELSVSSQVESLSRTPVLQSAWK
B.abortusBv3_Tulya         YGHGIIDNWLQPIRDIAQANQAELDTIENTQDRLDRLCELSVSSQVESLSRTPVLQSAWK
B.suisBv5_513              YGHGIIDNWLQPIRDIAQANQAELDTIENTQDRLDRLCELSVSSQVESLSRTPVLQSAWK
B.sp_NVSL_07-0026          YGHGIIDNWLQPIRDIAQANQAELDTIENTQDRLDRLCELSVSSQVESLSRTPVLQSAWK
B.suisBv2_ATCC23445        YGHGIIDNWLQPIRDIAQANQAELDTIENTQDRLDRLCELGVSSQVESLSRTPVLQSAWK
B.microti_CCM4915          YGHGIIDNWLQPIRDIAQANQAELDTIENTQDRLDRLCELSVSSQVESLSRTPVLQSAWK
B.suisBv4_40               YGHGIIDNWLQPIRDIAQANQAELDTIENTQDRLDRLCELSVSSQVESLSRTPVLQSAWK
                           ****************************************.*******************

B.abortusBv1_2308A         DGKDIIVHGWMYNLKDGLLRDIGCDCTRNALQFACQPAE
B.abortusBv5_B3196         DGKDIIVYGWMYNLKDGLLRDIGCDCTRNALQFACQPAE
B.spF5/99                  DGKDIIVHGWMYNLKDGLLRDIGCDCTRNALQFACQPAE
B.melitensisBv1_16M        DGKDIIVHGWMYNLKDGLLRDIGCDCTRNALQFACQPAE
B.melitensisBv2_63/9       DGKDIIVHGWMYNLKDGLLRDIGCDCTRNALQFACQPAE
B.suisBv1_1330             DGKDIIVHGWMYNLKDGLLRDIGCDCTRNALQFACQPAE
B.ceti_M13/05/1            DGKDIIVHGWMYNLKDGLLRDIGCDCTRNALQFACQPAE
B.abortusBv3_Tulya         DGKDIIVHGWMFNLKDGLLRDIGCDCTRNALQFACQPAE
B.suisBv5_513              DGKDIIVHGWMYNLKDGLLRDIGCDCTRNALQFACQPAE
B.sp_NVSL_07-0026          DGKDIIVHGWMYNLKDGLLRDIGCDCTRNALQFACQPAE
B.suisBv2_ATCC23445        DGKDIIVHGWMYNLKDGLLRDIGCDCTRNALQFACQPAE
B.microti_CCM4915          DGKDIIVHGWMYNLKDGLLRDIGCDCTRNALQFACQPAE
B.suisBv4_40               DGKDIIVHGWMYNLKDGLLRDIGCDCTRNALQFACQPAE
                           *******:***:***************************
```
